# Hydrogen Sulfide Deficiency and Therapeutic Targeting in Cardiometabolic HFpEF: Evidence for Synergistic Benefit with GLP-1/Glucagon Agonism

**DOI:** 10.1101/2024.09.16.613349

**Authors:** Jake E. Doiron, Mahmoud H. Elbatreek, Huijing Xia, Xiaoman Yu, Natalie D. Gehred, Tatiana Gromova, Jingshu Chen, Ian H. Driver, Naoto Muraoka, Martin Jensen, Smitha Shambhu, W.H. Wilson Tang, Kyle B. LaPenna, Thomas E. Sharp, Traci T. Goodchild, Ming Xian, Shi Xu, Heather Quiriarte, Timothy D. Allerton, Alexia Zagouras, Jennifer Wilcox, Sanjiv J. Shah, Josef Pfeilschifter, Karl-Friedrich Beck, Thomas M. Vondriska, Zhen Li, David J. Lefer

**Affiliations:** Department of Cardiac Surgery, Smidt Heart Institute, Cedars-Sinai Medical Center, Los Angeles, CA; Department of Pharmacology and Cardiovascular Center, Louisiana State University Health Sciences Center, New Orleans, LA; Department of Anesthesiology, Medicine & Physiology, David Geffen School of Medicine at University of California, Los Angeles, CA; Gordian Biotechnology, South San Francisco, CA; Department of Cardiovascular Medicine, Heart, Vascular and Thoracic Institute, Cleveland Clinic, Cleveland, OH; Center of Microbiome and Human Health, Department of Cardiovascular and Metabolic Sciences, Lerner Research Institute, Cleveland Clinic, Cleveland, Cleveland, OH; Molecular Pharmacology and Physiology, University of South Florida, Tampa, FL; Department of Chemistry, Brown University, Providence, RI; Vascular Metabolism Laboratory, Pennington Biomedical Research Center, Baton Rouge, LA; Northwestern University Medicine, Feinberg School of Medicine, Chicago, IL; Institute of Pharmacology and Toxicology, Goethe University, Frankfurt am Main, Germany; Department of Pharmacology and Toxicology, Faculty of Pharmacy, Zagazig University, Zagazig, Egypt

**Author notes:** J.D. and M.E. contributed equally. **Address for correspondence:** David J. Lefer, Ph.D., Department of Cardiac Surgery, Smidt Heart Institute, Cedars-Sinai Medical Center, Los Angeles, CA 90048. **Tweet:** H_2_S deficiency is linked to HFpEF in humans and animal models. Reduced production by CSE and increased metabolism by SQR impair H_2_S bioavailability. H_2_S donor synergizes with GLP-1/glucagon agonist to ameliorate HFpEF. These findings suggest a promising therapeutic approach for HFpEF. #**LeferLab** #**HeartFailure** #**H_2_SResearch**.

**Keywords:** H_2_S, HFpEF, CSE, SQR, GLP-1, Survodutide

## Abstract

**Background:** Heart failure with preserved ejection fraction (HFpEF) is a significant public health concern with limited treatment options. Dysregulated nitric oxide-mediated signaling has been implicated in HFpEF pathophysiology, however, little is known about the role of endogenous hydrogen sulfide (H_2_S) in HFpEF.

**Objectives:** This study evaluated H_2_S bioavailability in patients and two animal models of cardiometabolic HFpEF and assessed the impact of H_2_S on HFpEF severity through alterations in endogenous H_2_S production and pharmacological supplementation. We also evaluated the effects of the H_2_S donor, diallyl trisulfide (DATS) in combination with the GLP-1/glucagon receptor agonist, survodutide, in HFpEF.

**Methods:** HFpEF patients and two rodent models of HFpEF (“two-hit” L-NAME + HFD mouse and ZSF1 obese rat) were evaluated for H_2_S bioavailability. Two cohorts of two-hit mice were investigated for changes in HFpEF pathophysiology: (1) endothelial cell cystathionine-γ-lyase (EC-CSE) knockout; (2) H_2_S donor, JK-1, supplementation. DATS and survodutide combination therapy was tested in ZSF1 obese rats.

**Results:** H_2_S levels were significantly reduced (i.e., 81%) in human HFpEF patients and in both preclinical HFpEF models. This depletion was associated with reduced CSE expression and activity, and increased SQR expression. Genetic knockout of H_2_S -generating enzyme, CSE, worsened HFpEF characteristics, including elevated E/e’ ratio and LVEDP, impaired aortic vasorelaxation and increased mortality. Pharmacologic H_2_S supplementation restored H_2_S bioavailability, improved diastolic function and attenuated cardiac fibrosis corroborating an improved HFpEF phenotype. DATS synergized with survodutide to attenuate obesity, improve diastolic function, exercise capacity, and reduce oxidative stress and cardiac fibrosis.

**Conclusions:** H_2_S deficiency is evident in HFpEF patients and conserved across multiple preclinical HFpEF models. Increasing H_2_S bioavailability improved cardiovascular function, while knockout of endogenous H_2_S production exacerbated HFpEF pathology and mortality. These results suggest H_2_S dysregulation contributes to HFpEF and increasing H_2_S bioavailability may represent a novel therapeutic strategy for HFpEF. Furthermore, our data demonstrate that combining H_2_S supplementation with GLP-1/glucagon receptor agonist may provide synergistic benefits in improving HFpEF outcomes.

**Highlights:**

- H_2_S deficiency is evident in both human HFpEF patients and two clinically relevant models.
- Reduced H_2_S production by CSE and increased metabolism by SQR impair H_2_S bioavailability in HFpEF.
- Pharmacological H_2_S supplementation improves diastolic function and reduces cardiac fibrosis in HFpEF models.
- Targeting H_2_S dysregulation presents a novel therapeutic strategy for managing HFpEF.
- H_2_S synergizes with GLP-1/glucagon agonist and ameliorates HFpEF

## INTRODUCTION

Heart failure with preserved ejection (HFpEF) fraction is a significant public health concern. The increasing prevalence of HFpEF risk factors has resulted in HFpEF diagnoses accounting for 50% of all new heart failure diagnoses, while the subsequent pathophysiology leaves HFpEF patients subject to a 23% 30-day hospital readmission rate and 50-75% 5-year mortality rate^1-4^. In order to effectively treat HFpEF it is crucial that we further our understanding of the cardiovascular, and systemic drivers of this devastating CV disease. The influence of the physiologic gaseous mediator, nitric oxide (NO), has been extensively investigated in both preclinical and clinical studies^5,6^. While the precise role of NO signaling and nitrosative stress in HFpEF is still being elucidated, little is known about other physiological gaseous molecules, particularly hydrogen sulfide (H_2_S).

Since its discovery as an endogenously produced and biologically essential molecule in 1996, H_2_S has garnered significant attention for its role in maintaining whole-body and cardiovascular homeostasis^7,8^. H_2_S biosynthesis is regulated through three key enzymes, cystathionine-β-synthase (CBS), cystathionine-γ-lyase (CSE) and 3-mercaptopyruvate sulfurtransferase (3-MST)^9-11^. The major catabolic pathway for H_2_S breakdown occurs in the mitochondria under the control of sulfide quinone oxidoreductase (SQR)^12^. Perturbations in H_2_S bioavailability have been implicated in numerous cardiovascular pathologies, including atherosclerosis, hypertension, myocardial infarction, and heart failure with reduced ejection fraction (HFrEF)^13-16^. Investigation of H_2_S in these pathologies has revealed that H_2_S exerts potent antioxidant, anti-inflammatory, metabolic regulatory, and mitochondrial preservation actions^17-21^. Additionally, H_2_S has significant interplay with NO, whereby H_2_S augments cardioprotective endothelial nitric oxide synthase (eNOS) signaling^22^. Considering that HFpEF is characterized by derangements in metabolism, inflammation, oxidative stress and NO signaling, there is strong rationale for elucidation of the potential role of alterations in H_2_S signaling in HFpEF.

Of the HFpEF phenotypes, cardiometabolic HFpEF is of increasing importance given its reputation as the most prevalent HFpEF phenotype in a syndrome growing in incidence and diagnoses^1,23^. Cardiometabolic HFpEF comprises an estimated 33% of the HFpEF patient population and is most frequently comorbid with obesity, diabetes, hypertension, chronic inflammation, and hepatic injury^24,25^. Interestingly, H_2_S deficiency has been associated with these independent comorbidities and replenishment of H_2_S has been shown to improve the respective pathologies^10,18,26-30^.

Given the complexity of HFpEF, a multi-pronged approach targeting various underlying mechanisms is likely more effective than single-drug treatments. Glucagon-like peptide-1 receptor agonists (GLP-1RAs) are already approved for obesity and diabetes and are recommended by professional and medical societies for mitigation of the cardiovascular risk in patients with type 2 diabetes^31^. These drugs are categorized by the receptors they activate. Single GLP-1RAs activate only the GLP-1 receptor (e.g., liraglutide, dulaglutide, semaglutide), while dual GLP-1 and GIP RAs activate both GLP-1 and glucose-dependent insulinotropic polypeptide (GIP) receptors (e.g., tirzepatide). Both have shown promise in recent HFpEF clinical trials^32,33^. Emerging classes include dual GLP-1 and glucagon RAs, which activate both GLP-1 and glucagon receptors (e.g., mazdutide, survodutide) and are currently under development for obesity, diabetes and have shown very powerful effects against metabolic liver disease^34^. Finally, triple GLP-1, GIP and glucagon RAs activate all three receptors (e.g., retatrutide), demonstrating strong anti-obesity effects compared to single and dual agonists^35,36^. While the potential of the later two classes in HFpEF is intriguing, their impact remains to be determined.

In the present study, we sought to evaluate alterations in H_2_S bioavailability and the role of H_2_S in HFpEF using multiple clinically relevant models of cardiometabolic HFpEF. We also investigated the potential beneficial actions of H_2_S therapy in the form of H_2_S donors in the setting of HFpEF. This study also aimed to explore Survodutide’s therapeutic benefits in HFpEF and investigate if its effects are enhanced when combined with an H_2_S donor.

## METHODS

### Human Plasma Samples

Plasma samples were collected from either ambulatory patients with HFpEF prospectively enrolled in a heart failure outpatient clinic (with signs and symptoms or heart failure and echocardiogram showing left ventricular ejection fraction ≥50%) or age- and sex-matched healthy participants recruited from the community without existing cardiovascular diseases (confirmed by echocardiography, pulmonary function testing, and cardiac biomarkers) under protocols approved by the Institutional Review Board (IRB# 06-805 and #10-727, respectively) at the Cleveland Clinic. All participants provided written informed consent to participate. Baseline characteristics of all participants are shown in ***Table 1***.

**Table 1.**
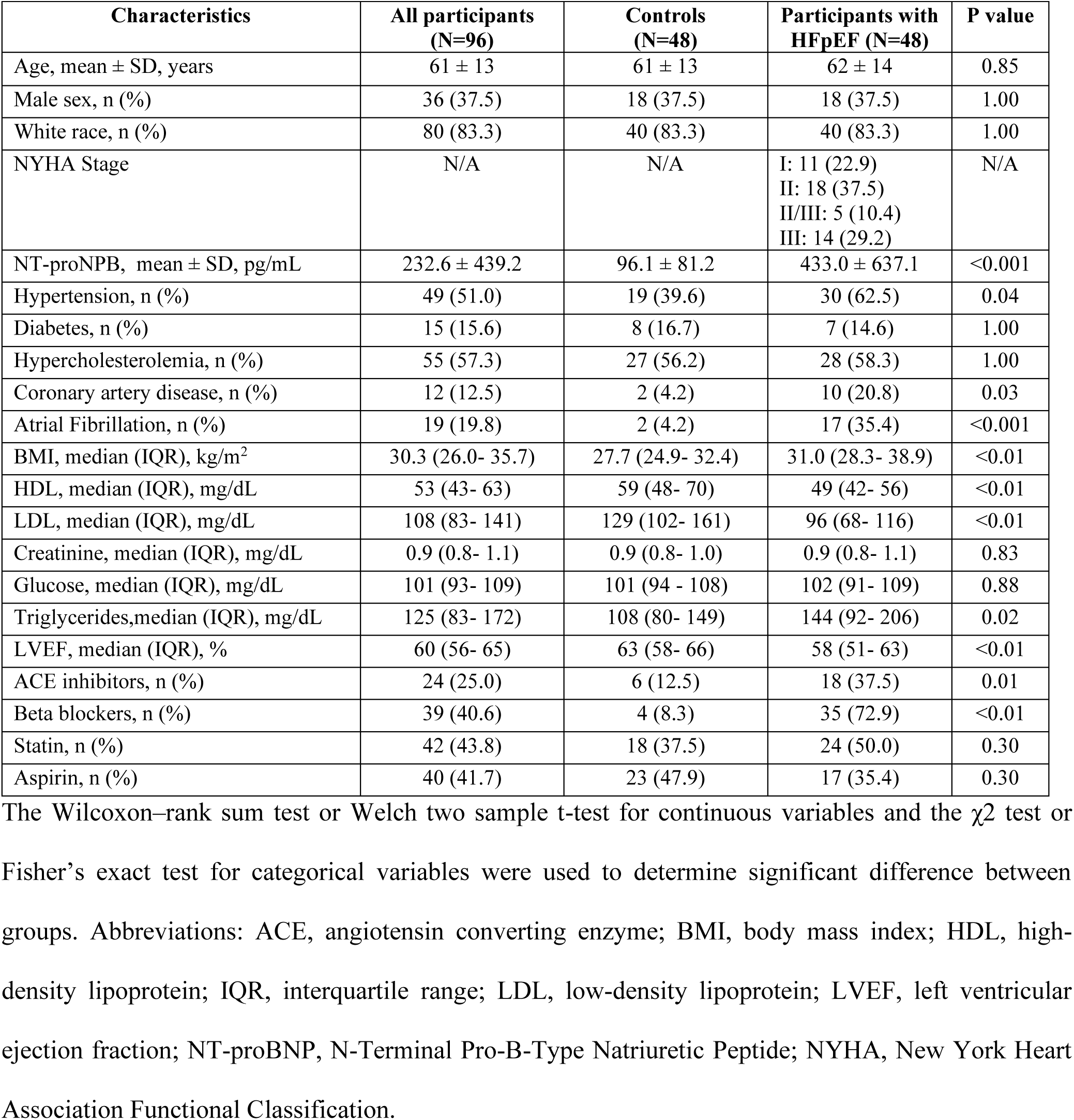
Baseline Characteristics of participants with HFpEF vs. Healthy Controls.

| Characteristics | All participants<br>(N=96) | Controls<br>(N=48) | Participants with<br>HFpEF (N=48) | P value |
| --- | --- | --- | --- | --- |
| Age, mean ± SD, years | 61 ± 13 | 61 ± 13 | 62 ± 14 | 0.85 |
| Male sex, n (%) | 36 (37.5) | 18 (37.5) | 18 (37.5) | 1.00 |
| White race, n (%) | 80 (83.3) | 40 (83.3) | 40 (83.3) | 1.00 |
| NYHA Stage | N/A | N/A | I: 11 (22.9)<br>II: 18 (37.5)<br>II/III: 5 (10.4)<br>III: 14 (29.2) | N/A |
| NT-proNPB, mean ± SD, pg/mL | 232.6 ± 439.2 | 96.1 ± 81.2 | 433.0 ± 637.1 | <0.001 |
| Hypertension, n (%) | 49 (51.0) | 19 (39.6) | 30 (62.5) | 0.04 |
| Diabetes, n (%) | 15 (15.6) | 8 (16.7) | 7 (14.6) | 1.00 |
| Hypercholesterolemia, n (%) | 55 (57.3) | 27 (56.2) | 28 (58.3) | 1.00 |
| Coronary artery disease, n (%) | 12 (12.5) | 2 (4.2) | 10 (20.8) | 0.03 |
| Atrial Fibrillation, n (%) | 19 (19.8) | 2 (4.2) | 17 (35.4) | <0.001 |
| BMI, median (IQR), kg/m<sup>2</sup> | 30.3 (26.0- 35.7) | 27.7 (24.9- 32.4) | 31.0 (28.3- 38.9) | <0.01 |
| HDL, median (IQR), mg/dL | 53 (43- 63) | 59 (48- 70) | 49 (42- 56) | <0.01 |
| LDL, median (IQR), mg/dL | 108 (83- 141) | 129 (102- 161) | 96 (68- 116) | <0.01 |
| Creatinine, median (IQR), mg/dL | 0.9 (0.8- 1.1) | 0.9 (0.8- 1.0) | 0.9 (0.8- 1.1) | 0.83 |
| Glucose, median (IQR), mg/dL | 101 (93- 109) | 101 (94 - 108) | 102 (91- 109) | 0.88 |
| Triglycerides, median (IQR), mg/dL | 125 (83- 172) | 108 (80- 149) | 144 (92- 206) | 0.02 |
| LVEF, median (IQR), % | 60 (56- 65) | 63 (58- 66) | 58 (51- 63) | <0.01 |
| ACE inhibitors, n (%) | 24 (25.0) | 6 (12.5) | 18 (37.5) | 0.01 |
| Beta blockers, n (%) | 39 (40.6) | 4 (8.3) | 35 (72.9) | <0.01 |
| Statin, n (%) | 42 (43.8) | 18 (37.5) | 24 (50.0) | 0.30 |
| Aspirin, n (%) | 40 (41.7) | 23 (47.9) | 17 (35.4) | 0.30 |

The Wilcoxon–rank sum test or Welch two sample t-test for continuous variables and the $\chi^2$ test or Fisher’s exact test for categorical variables were used to determine significant difference between groups. Abbreviations: ACE, angiotensin converting enzyme; BMI, body mass index; HDL, high-density lipoprotein; IQR, interquartile range; LDL, low-density lipoprotein; LVEF, left ventricular ejection fraction; NT-proBNP, N-Terminal Pro-B-Type Natriuretic Peptide; NYHA, New York Heart Association Functional Classification.

### Measurement of Plasma hs-CRP

The levels of hsCRP (#EA101010, ORIGENE, Rockville, MD) in plasma samples from control of HFpEF patients were measured with commercially available ELISA assay kits according to manufacturer’s instructions.

### Experimental Animal Models of HFpEF

#### “Two-hit” Mouse Model of HFpEF

Male C57BL/6N (Charles Rivers Laboratories, Wilmington, MA, USA) mice were purchased at 8 weeks of age and allowed to acclimatize for one week prior to study enrollment. Starting at 9 weeks of age, male C57BL/6N were treated with either L-N^G^-nitro arginine methyl ester (L-NAME, Enzo Biochem, Farmingdale, NY, USA) in the drinking water (0.5 gram/L) and 60% kcal high fat diet (HFD, D12492, Research Diets, New Brunswick, NJ, USA) to induce HFpEF (n = 7-10 per group), or normal drinking water and normal diet (Teklad 2019s, Inotiv, Chicago, IL, USA) (n = 7-10 per group). C57BL/6N mice were maintained on HFpEF treatment for 0, 5 or 10 weeks prior to physiological characterization. Similarly, male endothelial cell-cystathionine-γ-lyase (EC-CSE) knockout (KO) with constitutive Cre expression restricted to the endothelium, EC-CSE transgenic (Tg) and age-matched littermate controls were generated as described previously^37,38^. At 9 weeks of age, animals were enrolled to L-NAME and HFD treatment for 18 weeks. Lastly, for the H_2_S treatment intervention with H_2_S donor, JK-1, male C57BL/6N were enrolled at 9 weeks of age and began L-NAME and HFD treatment. At 5 weeks of L-NAME and HFD, mice were further randomized to either receive JK-1 (100µg/kg, b.i.d, i.p.) or vehicle (saline, b.i.d, i.p.). Group sizes of n = 7-10 were determined using a power and sample analysis with the significance level at 5% and power at 80%.

#### ZSF1 Obese Rat Model of Cardiometabolic HFpEF

Wistar Kyoto and ZSF1 obese rats (Charles River Laboratories, Wilmington, MA, USA) were purchased at 8 weeks of age to acclimatize for at least 2 weeks prior to study enrollment. Group sizes of n = 6 - 8 were determined using a power and sample analysis with the significance level at 5% and power at 80%. Mice and rats were housed at LSUHSC and CSMC in a temperature controlled and 12-hour light/dark cycle. All studies were LSUHSC and CSMC IACUC (Institutional Animal Care and Use Committee) approved and received care in LSUHSC animal care according to AALAC guidelines.

A cohort of male ZSF1 obese rats were subjected to an 8-week treatment regimen starting at 18 weeks of age. Animals were randomly assigned to three groups: vehicle (saline, n=6), Survodutide (MedChemExpress LLC, USA, 30 nmol/kg, subcutaneously, biweekly, n=8), or a combination of Survodutide and DATS (Cayman Chemical, USA, 150 µM/kg, intraperitoneally, daily, n=8).

### Non-Fasting Blood Glucose

A blood drop from the rat tail was used to measure non-fasting blood glucose with an Accu-Check Guide glucometer (Roche, Indianapolis, IN, USA).

### Hydrogen Sulfide Donors

JK-1 was synthesized by Dr. Ming Xian et al. as previously reported^39^. The dosage of JK-1 (100 µg/kg, b.i.d) was determined based on our previous studies of JK-1 in murine models of CV disease ^40-42^. DATS was prepared as described previously^43^.

### Exercise Capacity Testing

Treadmill exercise capacity of either mice or rats was performed using an IITC Life Science 800 Series rodent specific treadmill (Woodland Hills, CA). For mice, animals were first allowed to acclimate to the treadmill environment for a period of 5 minutes. Following acclimation, the mice were subjected to a warm-up protocol on a flat plane starting at 1 meter per minute that began increasing by 1.1m/min until a final speed of 12 meters/min was achieved after 10 minutes and then subsequently maintained for an additional 5 minutes. After the 15-minute warm-up, the ramp was increased to a 30° incline and the exercise capacity of the mice were recorded during an experimental protocol starting at 12 meters/min that increased by 2 meters/min until a final speed of 18 meters/min was achieved and maintained until the mouse reached a state of exhaustion.

Rats were allowed to acclimate to the treadmill environment for a period of 5 minutes. After acclimation, a warm-up protocol on a flat plane was initiated consisting of a starting speed of 6m/min with a gradual increase of 1.5 meters/min until a final speed of 12 meters/min was achieved and maintained for an additional minute. After the 5-minute warm-up, the animal’s exercise capacity was recorded during an experimental run on a flat plane at a consistent speed of 12 meters/min until the animal reached a state of exhaustion. Exhaustion was determined as an animal refusing to run for greater than 5 seconds or an inability of the animal to reach the front of the treadmill for 20 seconds. Data are represented as both exercise distance and work (kg*m) utilizing the animal’s body weight immediately prior to exercise testing.

### Transthoracic Echocardiography

Echocardiography was performed utilizing a Vevo-2100 ultrasound system (Visual Sonics, Toronto, Canada) for mice and rats. Animals were shaved 18 hours prior to experimentation. Animals were induced at 3% isoflurane and maintained at 1-3% isoflurane for echocardiographic evaluation. For mice, recordings pertaining to diastolic function occurred at a target heart rate of >400 beats per minute (bpm) and >450 bpm for systolic measures. For rats, a target heart rate of >300 bpm was used for systolic and diastolic measures. Left ventricular ejection fraction (LVEF) was quantified using an M-mode image across the parasternal short-axis view. Early filling velocity (E), atrial filling velocity (A), and early diastolic tissue velocity (E’) were measured in the four-chamber apical view. Data are represented as averages of three consecutive measurements for each parameter.

### Systemic and Left Ventricular Hemodynamic Measurements

Left ventricular end-diastolic and systemic pressures were measured as previously described.^40^ In brief, at the study endpoint, animals were anesthetized at 3% isoflurane until unresponsive to stimuli. The right common carotid artery was further surgically isolated and exposed. Once exposed, the isoflurane was reduced to 1% while a 1.4Fr (for mice) or 1.6Fr (for rats) high-fidelity pressure catheter (Transonic, NY, USA) was inserted and measurements of systemic blood pressures were recorded. The catheter was then advanced into the left ventricular lumen for measurement of left ventricular end-diastolic pressure (LVEDP).

### *Ex Vivo* Aortic Ring Vascular Reactivity

Thoracic aortas of mice or rats were removed for vascular reactivity testing at sacrifice as previously described.^44^ In brief, thoracic aortic rings were first equilibrated in Krebs-Henseleit solution and provided a tension of 0.5 grams for 60 minutes to reach equilibration. The rings were then pretreated with phenylephrine for maximal constriction and then followed with challenges of titrated acetylcholine (10^-9^ to 10^-5^ M) and subsequently sodium nitroprusside (10^-9^ to 10^-5^ M) and measured relaxation as compared to phenylephrine maximal contraction. Data are reported as percent relaxation from the maximum contraction to phenylephrine.

### Hydrogen Sulfide and Sulfane Sulfur Measurements

Hydrogen sulfide and sulfane sulfur were measured in plasma and tissue samples utilizing a gas chromatography-sulfur chemiluminescence system (Agilent Technologies, Santa Clara, CA, USA) as previously described^19^.

### Real-Time PCR

Total RNA from tissues was extracted with TRIzol (Life Technologies Corporation, USA). First strand cDNAs were obtained with iScrip Reverse Transcription Supermix (Bio-Rad, USA). Quantitative real-time PCR was performed with SYBR Green qPCR Master Mix (Selleck Chemicals, USA) on CFX Duet (Bio-Rad, USA). 2^-ΔΔCt^ method was utilized to calculate relative expression to 18s/Tubb5. Primer sequences used in the current study are listed in **Supplemental Table 1**.

### Western blot

Immunoblot analysis was performed on samples lysed in RIPA lysis buffer (MedChemExpress, USA) with the addition of Phosphatase Inhibitor (Selleck Chemicals, USA) and Protease Inhibitor Cocktail Mini-Tablet (MedChemExpress, USA) using standard procedures. The protein concentration was determined with BCA Protein Assay Kits (Thermo, USA). Total protein was separated with SDS-PAGE and transferred onto PVDF membranes (Gen Hunter Corporation, USA). Western blot analysis was performed using commercially available antibodies: CSE Polyclonal Antibody (1:2,000, PA5-29725; Thermo, USA), Anti-SQRDL Antibody (1:1,000, 144-09256-20; Ray Biotech, USA). Total protein staining used as loading control was obtained utilizing No-Stain™ Protein Labeling Reagent (Invitrogen, USA). Where mentioned, proteins were assessed using ImageJ software to analyze the optical density of western blots normalized to loading control. Total protein quantification was performed using signals from the entire lane as loading control.

### Enzyme Activity Assay

Enzyme reactions to determine the production of H_2_S from CSE were performed as described previously^45^ with slight modification. Briefly, tissues were homogenized in buffer containing 100 mM potassium phosphate, pH 7.4. For enzyme reactions, 0.172 mL of homogenate was incubated with 0.028 mL of substrate mix (10 mM L-cysteine, 2 mM pyridoxal 5`′`-phosphate) in a sealed vial at 37°C for 30 minutes. After adding 0.400 ml of 1 M sodium citrate buffer (pH 6.0), the mixtures were incubated at 37°C for 15 min with shaking on a rotary shaker to facilitate a release of H_2_S gas from the aqueous phase.

After shaking, 0.1 mL of head-space gas was applied to a gas chromatograph as described above. For both reactions, the H_2_S concentration of each sample was calculated against a calibration curve of Na_2_S.

### Single Nuclei RNA-seq & Analysis

The left ventricle of the heart tissue was collected from WKY and ZSF1 Obese Rats at both 14 and 26 weeks of age. The collected tissue was snap-frozen in liquid nitrogen. Heart nuclei were isolated using a lysis buffer consisting 0.25M sucrose, 10mM Tris-Hcl pH 7.5, 25mM KCl, 5mM MgCl2, 45uM Actinomycin D, supplemented with 1X protease inhibitor (G6521,Promega), 0.4U/uL RNasin Ribonuclease Inhibitor (N2515, Promega), 0.2U/ul SuperaseIn (AM2694, ThermoFisher). Briefly, heart tissue samples were minced into smaller pieces with scissors in a 1ml lysis buffer. The minced tissue was homogenized in a dounce homogenizer on ice with 10 strokes of pestle A, followed by 10 strokes of pestle B. The homogenized heart tissue was filtered through a 40 μm cell strainer and centrifuged at 400 x g for 5 min at 4°C. The nuclei pellet was resuspended in 2% BSA in PBS supplemented with Protector RNase inhibitor (03335402001, Sigma-Aldrich) at 0.2 U/ul. Heart nuclei were stained with Sytox red (S34859, Thermo Fisher, Waltham, MA) and sorted and purified through fluorescence-activated cell sorting (FACS). 8000-1000 nuclei from each rat heart were processed using a 10X Genomics microfluidics chip to generate barcoded Gel Bead-In Emulsions according to manufacturer protocols. Indexed single-cell libraries were then created according to 10X Genomics specifications (Chromium Next GEM Single Cell 5ʹ v2.1-Dual Index Libraries). Samples were multiplexed and sequenced in pairs on an Illumina Novaseq X (Illumina, San Diego, CA). The sequenced data were processed into expression matrices with the Cell Ranger Single-cell software 9.0.0 (https://www.10xgenomics.com/support/software/cell-ranger/latest/release-notes/cr-release-notes).

FASTQ files were obtained from the base-call files from Novaseq X sequencer and subsequently aligned to the rat genome NCBI Rnor6.0, with a read length of 26 bp for cell barcode and unique molecule identifier (UMI) (read 1), 8 bp i7 index read (sample barcode), and 98 bp for actual RNA read (read 2). Each rat sample yielded approximately 300 M reads.

Anndata and Scanpy were used to load and preprocess h5ad files for each sample for import into Seurat. Scrublet was used to identify and remove putative doublets before AnnData objects were concatenated by week. Barcodes, features, matrix files, and metadata were extracted into a folder for import into R with Seurat’s Read10X function. 14-week and 26-week Seurat objects were created with CreateSeuratObject. Mitochondrial gene percentage was calculated for each cell in each Seurat object before removing cells containing greater than 5% mitochondrial reads. Mitochondrial genes and genes with fewer than 10 reads across all cells in each object were also removed. Lastly, any cells with fewer than 500 reads were removed. The 14- and 26-week Seurat objects were merged by timepoint and layers joined. Samples were split by week and log-normalized with NormalizeData. FindVariableFeatures was used to identify the top 2000 variable features per week and the features were centered and scaled with ScaleData. Principle Component Analysis (RunPCA function) was run on the split objects with default parameters before layers were integrated together (IntegrateLayers function) with method=RPCAIntegration. A shared nearest-neighbor graph was created with FindNeighbors (15 RPCA dimensions) and clustered with FindClusters (resolution=0.5). Clusters were assigned cell types based on known marker genes. To create the heatmap in **Figure 5M**, the integrated Seurat object was subset into separate objects by cell type and week. Seurat’s FindMarkers function was used to identify differentially expressed genes in ZSF1 Obese cardiomoyoctes at 14 and 26 weeks, with a Bonferroni-corrected p-value cutoff of 0.05. Results were intersected with a list of known H2S-related enzymes, and the average Log2Fold-Change (FC) was plotted with ggplot2’s geom_tile function. To create the boxpot in **Figure 5N**, Seurat’s AverageExpression was used with feature=Sqor in cardiomyocytes grouped by rat, week, and condition. Average expression per cell per sample was plotted with GraphPad Prism 10.4.

### Histology and Fibrosis

Cardiac, renal cortex, and hepatic samples were collected and preserved in a 10% zinc formalin buffered solution (CUNZF-5-G, Azer Scientific, Morgantown, PA, USA) for 48 hours, after which they were transferred to a storage solution of 0.01% sodium azide in PBS. The samples were then embedded in paraffin and sectioned into 5 μm thick cross-sections. These sections were stained using either hematoxylin and eosin or picrosirius red with fast green counterstaining. Frozen liver sections from the rats were fixed and stained with Oil Red O as previously described^46^.

Images were processed with QuPath software and analyzed using ImageJ to quantify cardiac interstitial and perivascular fibrosis, as well as renal interstitial, perivascular, and periglomerular fibrosis, along with hepatic lipid content. For each animal, a minimum of 10 areas, vessels, or glomeruli within the tissue section were analyzed. Initially, images were converted to 8-bit grayscale with an RGB stack, and the threshold was adjusted to quantify lipid droplet features and the percentage of fibrosis.

### Glucose tolerance test

An oral glucose tolerance test was performed on rats. Fasting blood glucose was measured after a 6-hour fast. Subsequently, an oral glucose bolus (2 g/kg) was administered, and blood glucose levels were monitored at 0, 15, 30, 60, and 120 minutes using an Accu-Chek Guide glucose meter (Roche Diagnostics).

### Plasma and hepatic lipid assays

Plasma and hepatic triglycerides (# MAK266) and cholesterol (# MAK043) were measured by commercially available kits (Millipore Sigma, USA) according to manufacturer’s protocols.

### ELISA

Plasma 8-isoprostane (# 516351 Cayman Chemical, USA) and 3-nitrotyrosine (# NBP2-66363, Novus Biologicals, USA) levels were measured by ELISA kits according to manufacturer’s instructions.

### Statistical Analysis

Data are presented using both mean ± SEM and box plots (median, minimum, and maximum). All statistical analysis were performed in a blinded manner using the Prism 6 and Prism 10 software (GraphPad, San Diego, California). Differences in data among 2 groups were compared using an unpaired student t-test. For comparisons amongst multiple, separate groups of animals enrolled within the same experimental study, multiple unpaired student *t* tests were utilized. For data involving animal cohorts that were continuously monitored and evaluated according to factors such as timepoints or concentrations, 2-way ANOVA analysis followed by a Bonferroni multiple comparison test was utilized. Survival data was acquired using a Kaplan-Meier survival curve analysis through Prism 6. A p value of <0.05 was considered statistically significant. The presented data may have different numbers of animals per group as only a subset of the mice or rats from each group were used for certain experiments. Additionally, exclusion of animals was carried out due to complications such as procedural failure in invasive hemodynamics measurement, limited amount of sample collected (i.e., plasma volume), and lack of participation in treadmill running. Prior to conducting statistical analysis, an outlier test was performed using a Grubbs test (α = 0.05) through GraphPad to identify and remove any outliers in the data set.

## RESULTS

### H_2_S is Depleted in Human HFpEF

To investigate the role of H_2_S in HFpEF, we collected plasma samples from patients with confirmed HFpEF (n=46) and a control group without HFpEF (n=45). The characteristics of the patient population are detailed in Table 1. Our analysis revealed an 81% reduction in circulating H_2_S levels in the HFpEF patients, accompanied by significant decreases in sulfane sulfur levels, a stable metabolite of gaseous H_2_S (***Figure 1A and 1B***). Given that cardiometabolic HFpEF is characterized by low-grade chronic inflammation and that H_2_S possesses anti-inflammatory effects^47,48^, we measured circulating hs-CRP levels. We observed elevated levels of the inflammatory marker hs-CRP in the HFpEF group (***Figure 1C***). We also investigated potential correlations between circulating H_2_S and sulfane sulfur levels with the clinical variables presented. However, no statistically significant correlations were found. This may be due to the relatively small sample size of our study, which limits our power to detect such correlations.

**Figure 1.**
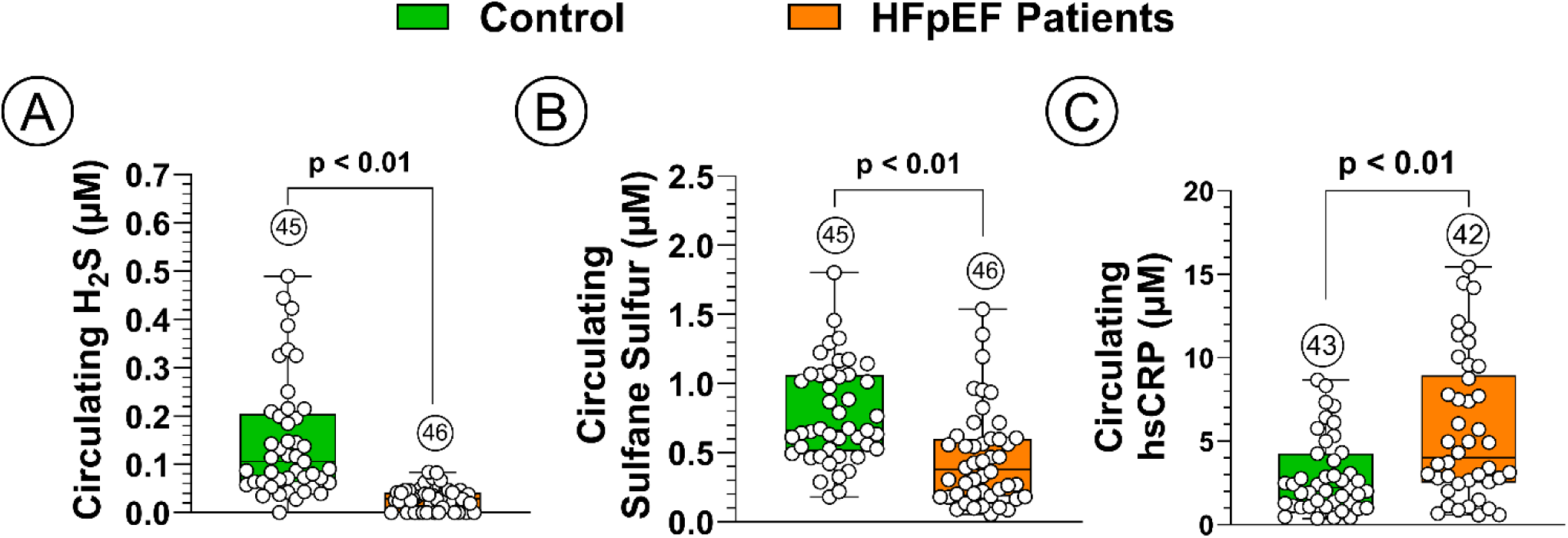
Reduced Circulating Hydrogen Sulfide in HFpEF Patients. *(**A**)* Circulating H_2_S (µM), *(**B**)* Circulating sulfane sulfur (µM), *(**C**)* Circulating hs-CRP (µM). Circled number inside bars indicate sample size. Data were analyzed with Student unpaired 2-tail *t* test. Data are presented as box plots (median, minimum, and maximum). hs-CRP, high-sensitivity C-reactive protein.

### H_2_S is Reduced in Mouse Two-Hit HFpEF Model Due to Decreased Production by CSE

We next employed a well-established “two-hit” cardiometabolic HFpEF mouse model (L-NAME + HFD)^5^, to assess H_2_S bioavailability and HFpEF phenotype after 0, 5, or 10 weeks of treatment as outlined in ***Supplemental Figure 1A***. We first characterized the HFpEF phenotype in this model. Mice with HFpEF exhibited increased body weights and elevated systolic and diastolic blood pressures (***Supplemental Figure 1B-1D***). While left ventricular ejection fraction (LVEF) remained within a preserved range, the E/e’ ratio increased throughout the treatment (***Supplemental Figure 1E and 1F***). Correspondingly, left ventricular catheterization revealed elevated left ventricular end-diastolic pressure (LVEDP), indicating increased myocardial stiffness and reduced compliance (***Supplemental Figure 1G***). Additionally, HFpEF mice displayed substantially diminished exercise capacity, as evidenced by reduced exercise distance (***Supplemental Figure 1H***). Endothelial dysfunction was also noted, with impaired aortic vasoreactivity to acetylcholine. Importantly, these changes in vascular reactivity were independent of altered responses to sodium nitroprusside, suggesting that the observed effects stem from compromised endothelial function (***Supplemental Figure 1I and 1J***). While our study demonstrates that both vascular dysfunction and cardiac abnormalities are present in the HFpEF model, establishing a definitive temporal relationship between these two physiological parameters requires further investigation. It is possible that endothelial dysfunction and impaired vascular reactivity may contribute to the development of cardiac abnormalities in HFpEF. For example, sustained increases in afterload due to impaired vascular compliance could contribute to the development of diastolic dysfunction and increased LVEDP. Further studies, such as longitudinal assessments of vascular function and cardiac parameters in HFpEF models, are necessary to fully elucidate the temporal and causal relationships between these two critical aspects of HFpEF pathophysiology.

At baseline, no differences were observed in circulating H_2_S; however, exposure to L-NAME and HFD resulted in a progressive decline in both circulating H_2_S and sulfane sulfur levels (***Figure 2A and 2B***). Similarly, cardiac H_2_S and sulfane sulfur levels decreased progressively throughout the two-hit regimen (***Figure 2C and 2D***). Although H_2_S levels was also reduced in control mice at later timepoints compared to baseline, we did not observe any changes in cardiac physiology as LV E/e’, remained stable over time. Potential contributing factors to this decline could include small sample size, age-related changes in H_2_S metabolism or subtle variations in experimental conditions across timepoints.

**Figure 2.**
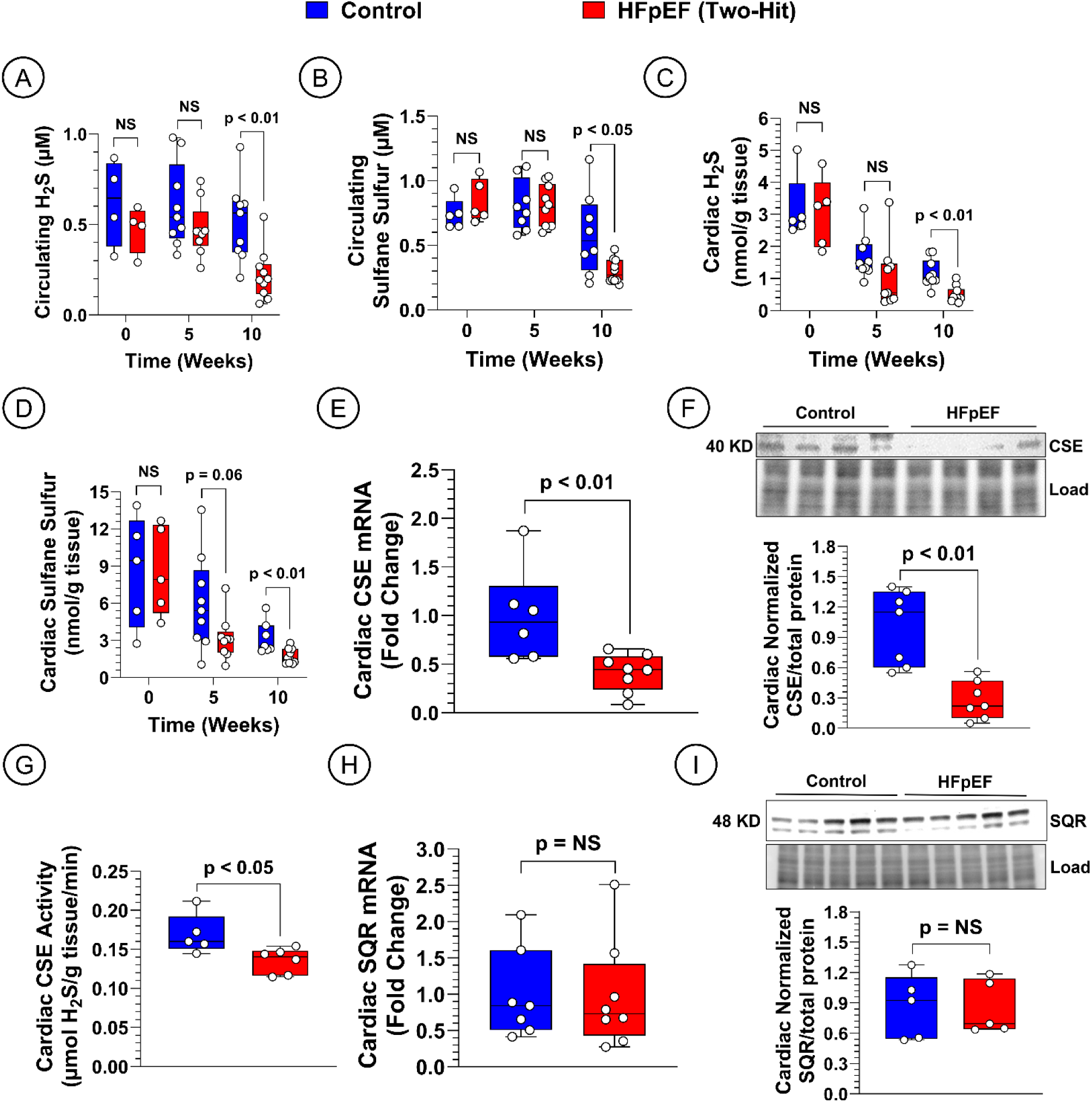
Circulating and Myocardial Hydrogen Sulfide Bioavailability in “Two-hit” Murine HFpEF Model. (***A***) Circulating H_2_S (µM), *(**B**)* Circulating sulfane sulfur (µM), *(**C**)* Myocardial H_2_S (nmol/g tissue), *(**D**)* Myocardial sulfane sulfur (nmol/g tissue), *(**E**)* Myocardial CSE gene expression, *(**F**)* Myocardial CSE protein expression expressed as fold change, *(**G**)* Myocardial CSE enzyme activity (nmol H_2_S/g tissue), *(**H**)* Myocardial SQR gene expression expressed as fold change, *(**I**)* Myocardial SQR protein expression expressed as fold change. Data in panels ***A***-***D*** were analyzed with multiple Student unpaired 2-tail *t* tests. Data in panels ***E***-***I*** were from tissues isolated after 10 weeks and analyzed with Student unpaired 2-tail *t* tests. Data are presented as box plots (median, minimum, and maximum). CSE, cystathionine γ-lyase; SQR, sulfide quinone oxidoreductase.

To investigate the underlying causes of H_2_S reduction in HFpEF, we examined the primary vascular H_2_S-producing enzyme, CSE, and the primary H_2_S-metabolizing enzyme, SQR^9,10^. In the hearts of HFpEF mice, both gene and protein expression, as well as enzyme activity of CSE, were significantly reduced (***Figure 2E-2G***), while SQR expression remained comparable to that in control hearts (***Figure 2H and 2I***)

Given the systemic nature of HFpEF, we also assessed H_2_S bioavailability in the liver and kidney which are the primary sources of systemic H_2_S production and regulate overall H_2_S bioavailability and metabolism. Indeed, liver is the highest CSE-expressing organ^12,49^. We observed significant reductions in hepatic H_2_S, along with decreased CSE gene and protein expression at the 10-week endpoint, although CSE activity did not differ from controls (***Supplemental Figure 2A-2D***). Notably, hepatic SQR gene expression was elevated in HFpEF, while protein levels remained unchanged (***Supplemental Figure 2E and 2F***). In the kidneys, no significant differences in H_2_S levels were noted between control and HFpEF models at the 10-week endpoint (***Supplemental Figure 2G***). However, renal CSE gene and protein expression, as well as enzyme activity, were significantly reduced in HFpEF, while SQR expression levels were similar to controls (***Supplemental Figure 2H-2L***).

These changes in systemic H_2_S bioavailability correlate with the worsening HFpEF phenotype. Our data demonstrate that the “two-hit” L-NAME + HFD murine model develops a progressively obese and hypertensive phenotype, accompanied by a systemic depletion of bioavailable H_2_S, primarily due to dysfunctional CSE in multiple organs.

### CSE Genetic Deficiency Exacerbates While Overexpression Attenuates HFpEF Phenotype

To further explore the potential causal relationship between CSE dysfunction and global H_2_S reduction, as well as to assess the impact of decreased H_2_S bioavailability on HFpEF severity, we investigated mice with genetic deletion or overexpression of endothelial cell cystathionine-γ-lyase (EC-CSE). EC-CSE knockout (KO) and transgenic (Tg) mice and their wild-type littermates underwent the same L-NAME and HFD HFpEF protocol. EC-CSE KO mice showed significantly reduced cardiac H_2_S levels and slightly reduced sulfane sulfur levels (***Figure 3A and 3B***). The lack of endothelial CSE resulted in elevated E/e’ ratios and LVEDP (***Figure 3C and 3D***). We also observed a significant impairment in exercise capacity in EC-CSE KO mice (***Figure 3E***). To assess vascular dysfunction in the context of CSE deficiency and HFpEF, we evaluated isolated thoracic aortic rings exposed to increasing concentrations of acetylcholine. Knockout animals demonstrated reduced vasorelaxation in response to acetylcholine, though responsiveness to sodium nitroprusside remained unchanged (***Figure 3F and 3G***). Notably, the genetic deficiency of EC-CSE was associated with increased mortality in our study, underscoring the critical role of H_2_S homeostasis in HFpEF (***Figure 3H***). Diastolic function and mortality from wild type control and EC-CSE KO treated with standard chow are presented in ***Supplemental Figure 3*.**

**Figure 3.**
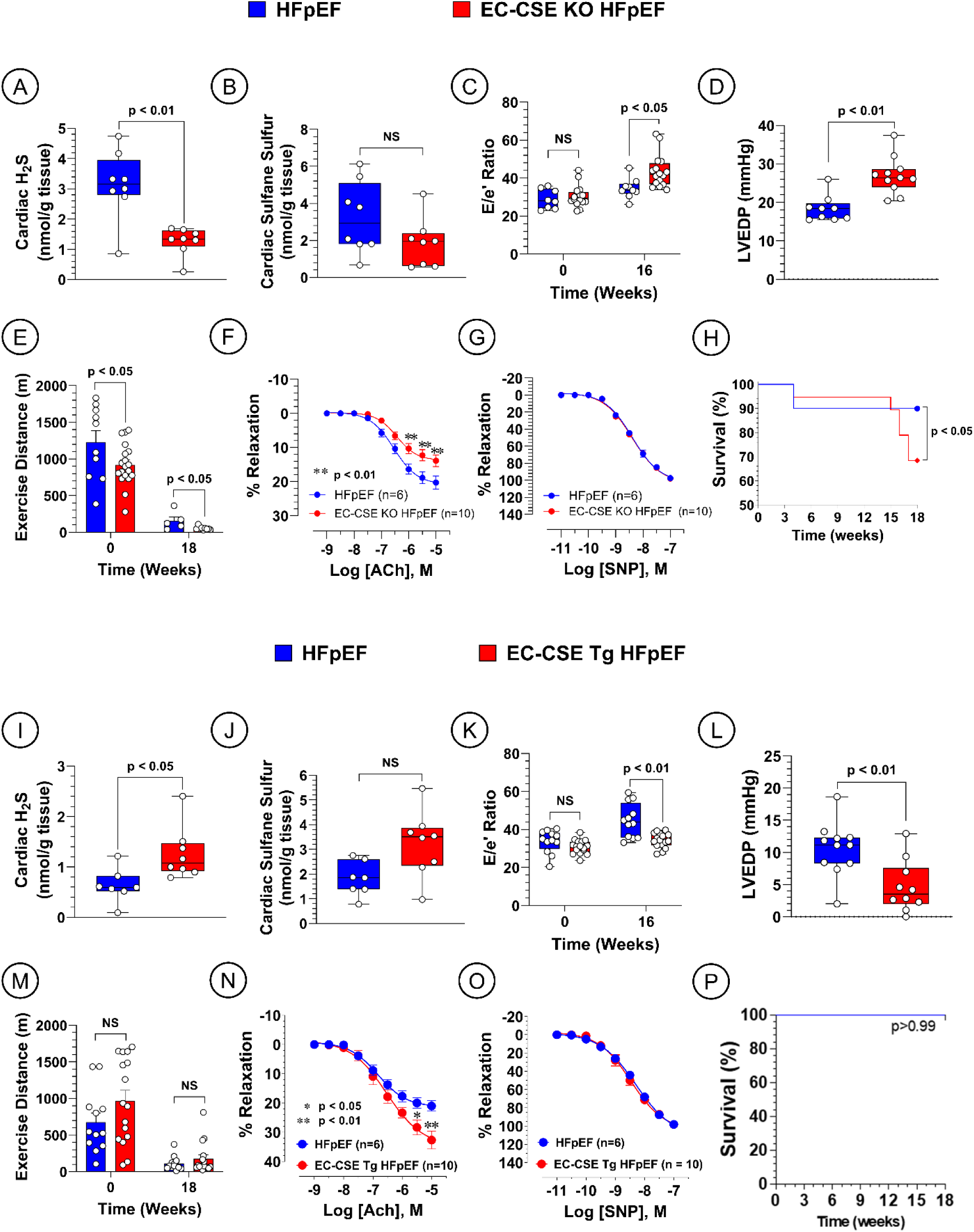
HFpEF in Endothelial Cell (EC) Cystathionine Gamma Lyase (CSE) Knockout and Transgenic Mice. *(**A**)* Myocardial H_2_S (nmol/g tissue), *(**B**)* Myocardial sulfane sulfur (nmol/g tissue), *(**C**)* Ratio of early mitral diastolic inflow velocity (E) and mitral annular early diastolic velocity (e’), (***D***) Left ventricular end diastolic pressure (LVEDP) in mmHg after 10 weeks, *(**E**)* Treadmill exercise distance in meters (m), *(**F**)* Aortic vascular reactivity to acetylcholine (ACh) after 10 weeks expressed as a percent of pre-contraction to norepinephrine, ***(G)*** Aortic vascular reactivity to sodium nitroprusside (SNP) after 10 weeks expressed as a percent of pre-contraction to norepinephrine, ***(H)*** Survival analysis. *(**I**)* Myocardial H_2_S (nmol/g tissue), *(**J**)* Myocardial sulfane sulfur (nmol/g tissue), *(**K**)* Ratio of early mitral diastolic inflow velocity (E) and mitral annular early diastolic velocity (e’), (***L***) Left ventricular end diastolic pressure (LVEDP) in mmHg after 10 weeks, *(**M**)* Treadmill exercise distance in meters (m), *(**N**)* Aortic vascular reactivity to acetylcholine (ACh) after 10 weeks expressed as a percent of pre-contraction to norepinephrine, ***(O)*** Aortic vascular reactivity to sodium nitroprusside (SNP) after 10 weeks expressed as a percent of pre-contraction to norepinephrine, ***(P)*** Survival analysis. Data in panels ***A, B, D, I, J and L*** were analyzed with Student unpaired 2-tail *t* test. Data in panels ***C, E, K* and *M*** were analyzed with multiple Student unpaired 2-tail *t* tests. Data in panels ***F, G, N* and *O*** were analyzed with ordinary 2-way ANOVA. Survival data in panels ***H and P*** was analyzed with Kaplan-Meier survival analysis. Data in panels ***A-D and I-L*** are presented as box plots (median, minimum, and maximum). Data in panels ***E-G and M-O*** are presented as mean ± SEM. * Echocardiography, # Treadmill exercise, ⚲ Aortic vascular reactivity, Invasive hemodynamics, H_2_S and sulfane sulfur measurements.

These findings are further supported by the studies in EC-CSE overexpressing transgenic mice, which demonstrate significantly higher H_2_S levels and provide protection against the development of HFpEF. Indeed, EC-CSE Tg mice exhibited higher levels of cardiac H_2_S and sulfane sulfur, albeit the latter was not statistically significant (***Figure 3I and 3J***). Transgenic mice had improved cardiac function as indicated by reduced E/e’ ratio and LVEDP (***Figure 3K and 3L)***. We did not observe any significant differences in exercise capacity between transgenic mice and controls (***Figure 3M)***, however the former showed a significant improvement in endothelium-dependent, but not endothelium-independent relaxation ***(Figure 3N and 3O).*** Remarkably, there was no mortality in both control and transgenic mice throughout the study (***Figure 3P)***.

### Exogenous H_2_S Administration Ameliorates Systemic HFpEF Dysfunction

From a translational perspective, we aimed to determine whether the H_2_S deficiency observed in HFpEF mice could be addressed with an exogenous pharmacological source of H_2_S. We enrolled a cohort of wild-type mice in the two-hit HFpEF protocol, where they received L-NAME and HFD for 5 weeks to induce HFpEF. After this period, mice were randomized to receive either a saline vehicle or the H_2_S donor JK-1 for an additional 5 weeks until the study endpoint (***Supplemental Figure 4A***). Mice treated with JK-1 exhibited significantly higher levels of circulating and cardiac sulfane sulfur, without notable increases in free H_2_S (***Supplemental Figure 4B-4E***). The short half-life of free H_2_S, combined with the pharmacokinetic profile of JK-1, likely accounts for the lack of elevated H_2_S levels; however, the increased sulfane sulfur indicates a significant enhancement in the physiological sulfur pool.

**Figure 4.**
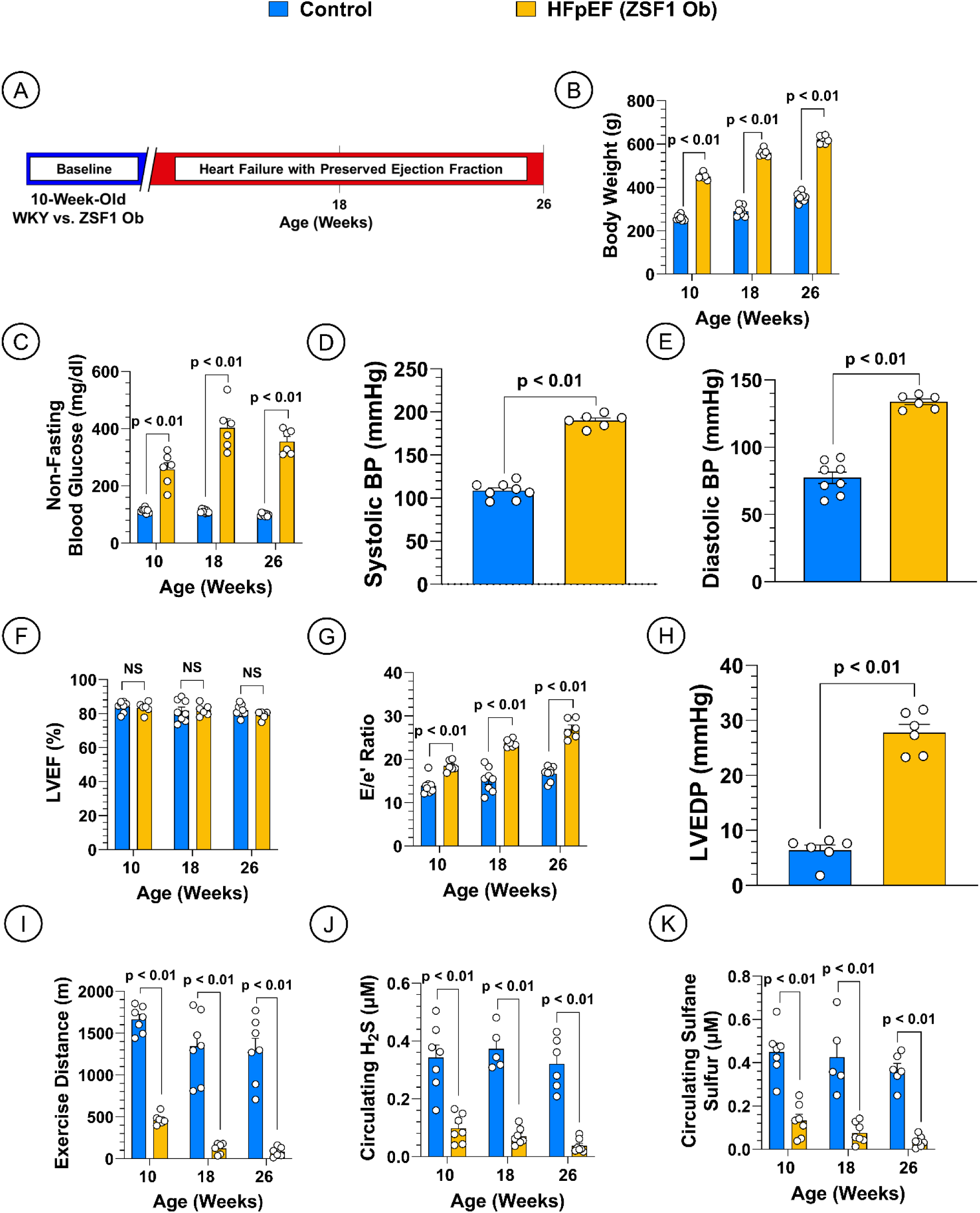
Reduced Circulating Hydrogen Sulfide Bioavailability and HFpEF Severity in the ZSF1 Obese Rat. *(**A**)* Study Timeline, (***B***) Body weight in grams (g), *(**C**)* Non-fasting blood glucose (mg/dl), (***D***) Systolic blood pressure (mmHg) at 26 weeks, *(**E**)* Diastolic blood pressure (mmHg) at 26 weeks, *(**F**)* Left ventricular ejection fraction (LVEF), *(**G**)* Ratio of early mitral diastolic inflow velocity (E) and mitral annular early diastolic velocity (e’), (***H***) Left ventricular end diastolic pressure (LVEDP) (mmHg) at 26 weeks, *(**I**)* Treadmill exercise distance expressed in meters (m), (***J***) Circulating H_2_S (µM), *(**K**)* Circulating sulfane sulfur (µM). Data in panels ***B, C, F, G, I-K*** were analyzed with multiple Student unpaired 2-tail *t* tests. Data in panels ***D, E and H*** were analyzed with Student unpaired 2-tail *t* test. Data are presented as mean ± SEM.

Exogenous H_2_S supplementation did not affect body weight, systolic blood pressure but resulted in a significant reduction in diastolic blood pressure (***Supplemental Figure 4F-4H***). Moreover, animals treated with JK-1 showed statistically significant decreases in both E/e’ and LVEDP (***Supplemental Figure 4I and 4J***). JK-1 treatment also significantly improved exercise capacity (***Supplemental Figure 4K***). These results are noteworthy, as HFpEF is characterized by severe exercise intolerance, and our findings suggest that H_2_S therapy enhanced exercise capacity even in the presence of an eNOS inhibitor, indicating that the benefits of H_2_S in HFpEF are likely independent of NO signaling^50^.

H_2_S has been shown to exert anti-fibrotic effects in models of myocardial infarction and HFrEF, prompting us to evaluate myocardial interstitial and perivascular fibrosis^40^. We observed significant reductions in both perivascular and interstitial fibrosis following JK-1 treatment (***Supplemental Figure 4L and 4M***). Additionally, our initial characterization of the L-NAME + HFD two-hit model indicated compromised hepatic H_2_S bioavailability. Hepatic H_2_S is crucial for liver health, and diminished hepatic H_2_S signaling has been linked to the development of non-alcoholic fatty liver disease (NAFLD), a condition recently proposed as a predictor of HFpEF diagnosis^26,51^. Notably, we found substantial reductions in hepatic lipid content in HFpEF mice treated with JK-1 for 5 weeks, further supporting the potential benefits of H_2_S in this systemic condition (***Supplemental Figures 4N***).

### Altered H_2_S Production by CSE and Metabolism by SQR Underpins Cardiometabolic HFpEF in ZSF1 Obese Rats

To further validate our earlier findings in humans and mice, we investigated H_2_S bioavailability in a clinically relevant model of cardiometabolic HFpEF using ZSF1 obese (Ob) rats^52,53^. For this study, we enrolled ZSF1 Ob rats and normotensive, non-obese, non-diabetic control Wistar Kyoto (WKY) rats. We comprehensively characterized the cardiometabolic HFpEF phenotype in ZSF1 Ob rats, monitoring HFpEF progression from week 10 to week 26 of age (***Figure 4A***). Key cardiometabolic parameters, including body weight, blood glucose levels, systolic and diastolic blood pressures, E/e’ ratio, and LVEDP, were significantly elevated in ZSF1 Ob rats compared to WKY controls, while LVEF was preserved, and exercise capacity was impaired (***Figure 4B-4I***). Notably, circulating H_2_S and sulfane sulfur levels were dramatically reduced as early as 10 weeks of age in the ZSF1 obese rats (***Figure 4J and 4K***). The reduction in circulating H_2_S and sulfane sulfur in the ZSF1 obese rat is very similar to that observed in HFpEF patients (i.e., ∼80-90%).

We also examined CSE and SQR expression in the heart, liver, and kidneys. While cardiac CSE gene expression did not differ between WKY and ZSF1 Ob rats, CSE protein levels were lower in the ZSF1 Ob group. Conversely, both SQR gene and protein expressions were elevated in these rats (***Figure 5A-5D***). In the liver, we observed significant reductions in both CSE gene and protein expression, with unchanged SQR gene expression but increased SQR protein levels in ZSF1 Ob rats (***Figure 5E-5H***). In the kidneys, CSE expression was increased, while SQR expression was decreased at both the gene and protein levels (***Figure 5I-5L***). Interestingly, single cell RNA sequencing revealed increased expression of SQR in the cardiomyocytes of ZSF1 Ob rats in both early HFpEF (14 weeks old) and late HFpEF (26 weeks old) (***Figure 5M and 5N***). This analysis also showed that both the H_2_S generator, CBS, and the H_2_S metabolizer, thiosulfate sulfurtransferase (TST), are elevated in the cardiomyocytes only in early HFpEF (***Figure 5M)***.

**Figure 5.**
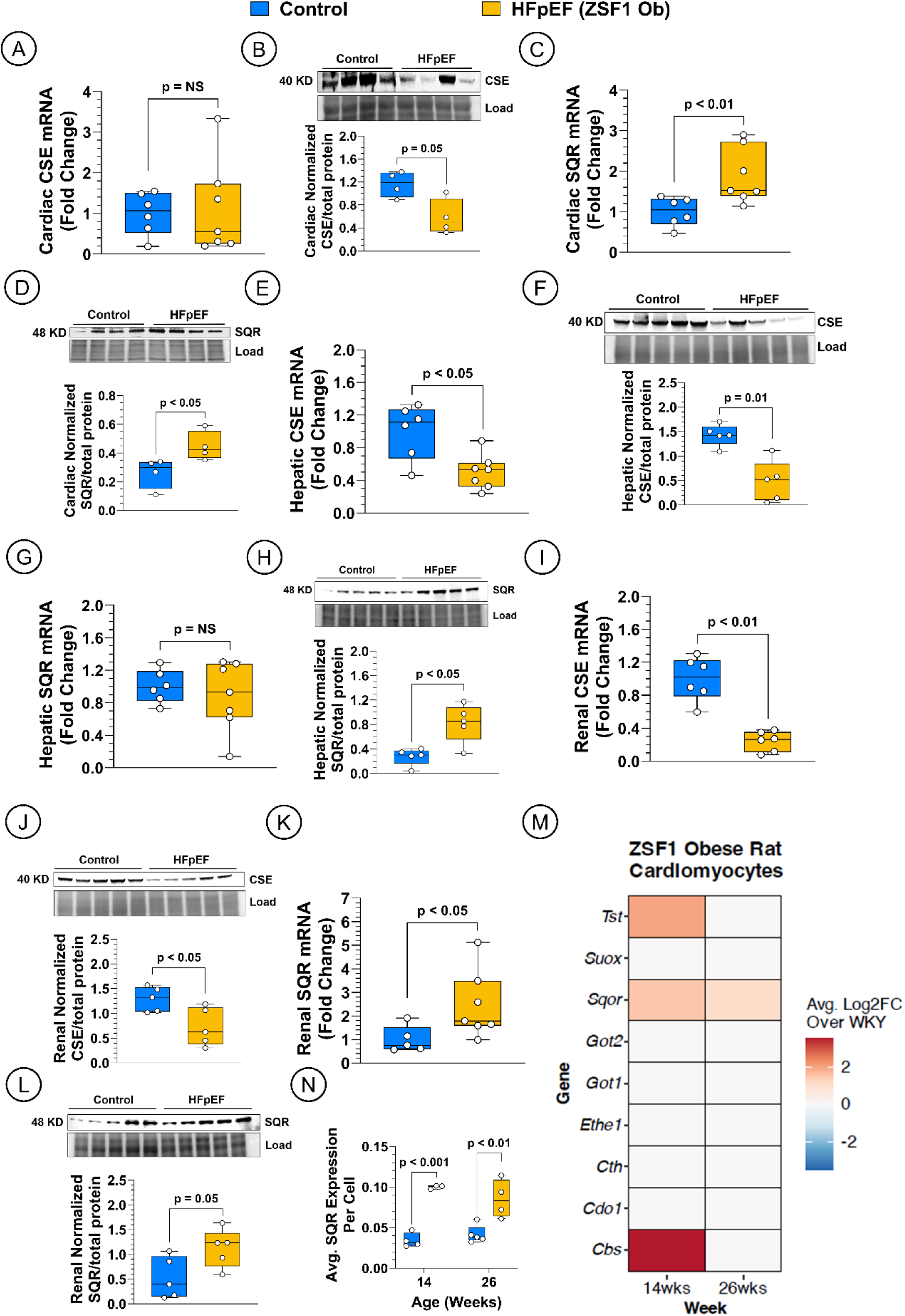
Reduced Expression of CSE and Increased Expression of SQR in the 26-Week-Old ZSF1 Obese Rat Tissues. *(**A**)* Myocardial CSE gene expression, (***B***) Myocardial CSE protein expression, *(**C**)* Myocardial SQR gene expression, (***D***) Myocardial SQR protein expression, *(**E**)* Hepatic CSE gene expression, (***F***) Hepatic CSE protein expression, *(**G**)* Hepatic SQR gene expression, ***(H)*** Hepatic SQR protein expression, *(**I**)* Renal CSE gene expression, (***J***) Renal CSE protein expression, *(**K**)* Renal SQR gene expression, ***(L)*** Renal SQR protein expression, *(**M**)* Heat map showing the expression of H_2_S related genes in rat cardiomyocytes, *(**N**)* Average SQR expression per cell. Data in ***A-L*** were analyzed with Student unpaired 2-tail *t* test. Data in ***N*** was analyzed with Two-Way ANOVA. Data are presented as box plots (median, minimum, and maximum). CSE, cystathionine γ-lyase; SQR, sulfide quinone oxidoreductase.

In the heart, interstitial and perivascular fibrosis were pronounced in ZSF1 Ob rats **(*Figure 6A and Supplemental Figure 5A***), and H_2_S levels were reduced despite unchanged CSE enzyme activity (***Figure 6B and 6C***). In the liver, lipid content was markedly increased, accompanied by substantial reductions in both H_2_S levels and CSE enzyme activity (***Figure 6D-6F***). The kidneys of ZSF1 Ob rats also exhibited significant perivascular, interstitial, and periglomerular fibrosis (***Figure 6G and Supplemental Figure 5B and 5C***), with reduced H_2_S levels and CSE enzyme activity observed by week 26 (***Figure 6H and 6I***).

**Figure 6.**
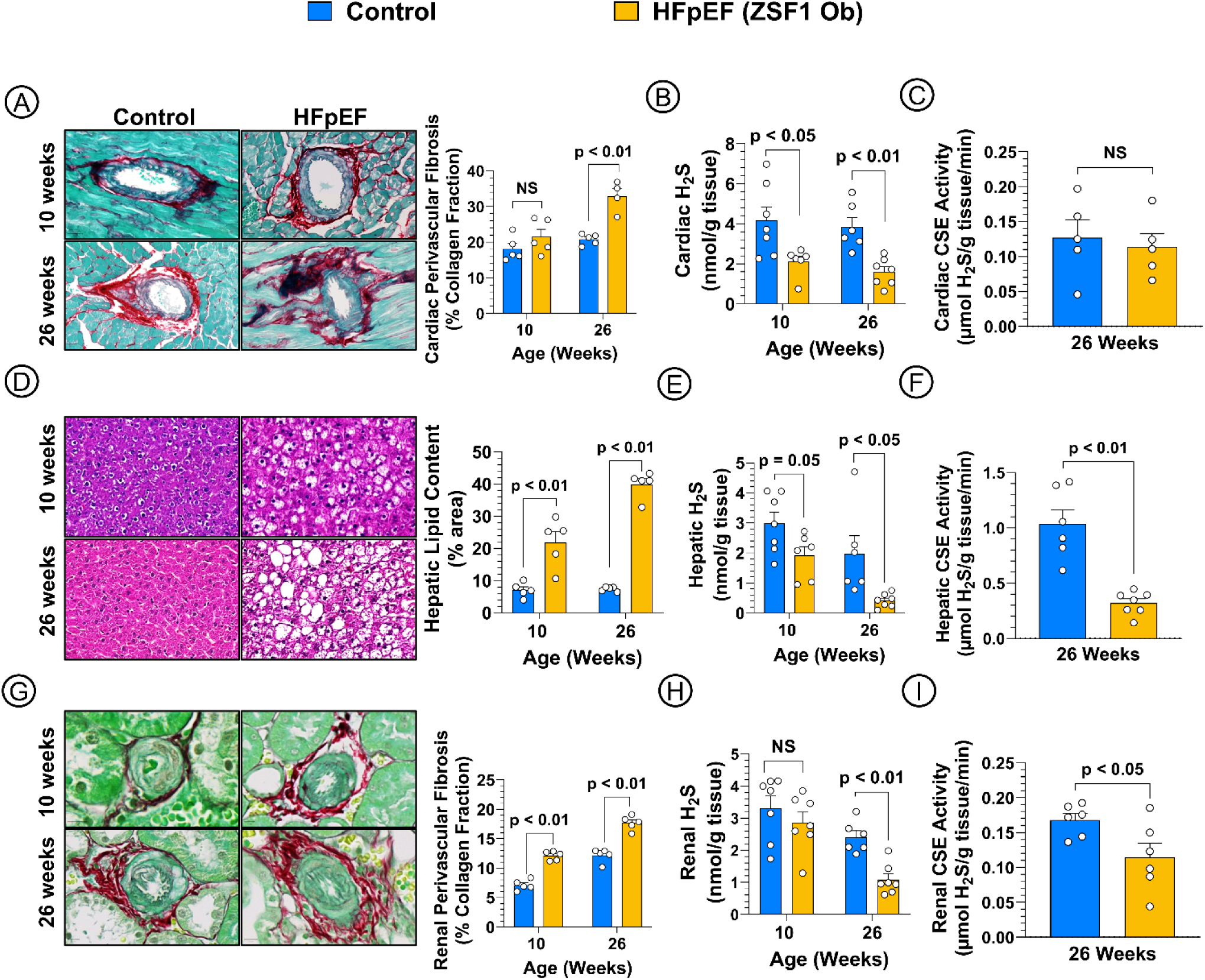
Histopathological Changes and Reduced Hydrogen Sulfide and CSE Enzyme Activity in the ZSF1 Obese Rats. ***(A)*** Representative images of cardiac perivascular fibrosis expressed as % collagen fraction at 10 and 26 weeks of age. Tissue stained with Masson’s Trichrome and fast green counterstaining and respective quantification, (***B***) Myocardial H_2_S (nmol/g tissue), *(**C**)* Myocardial CSE enzyme activity (µmol H_2_S/g tissue/min), *(**D**)* Representative images of hepatic lipid accumulation (% area) stained with hematoxylin and eosin and respective quantification, *(**E**)* Hepatic H_2_S (nmol/g tissue), (***F***) Hepatic CSE enzyme activity (µmol/g tissue/min), (***G***) Representative images of renal perivascular fibrosis (% collagen fraction) stained with Masson’s Trichrome and fast green counterstaining and respective quantification*, (**H**)* Renal H_2_S (nmol/g tissue), (***I***) Renal CSE enzyme activity (µmol/g tissue/min). Data in panels ***A, B, D, E, G and H*** were analyzed with multiple Student unpaired 2-tail *t* tests. Data in panels ***C, F and I*** were analyzed with Student unpaired 2-tail *t* test. Data are presented as mean ± SEM. CSE, cystathionine γ-lyase.

These findings further confirm global H_2_S deficiency as a consistent feature across multiple HFpEF models.

### H_2_S Synergizes with the GLP-1/Glucagon Receptor Survodutide in Ameliorating Cardiometabolic HFpEF in ZSF1 Obese Rats

To investigate a clinically relevant approach to treat HFpEF, we examined the therapeutic efficacy of Survodutide, a GLP-1/glucagon receptor agonist in combination with the well-characterized polysulfide H_2_S donor, DATS, in ZSF1 obese rats (***Figure 7A***). We confirmed that DATS treatment significantly increased circulating H_2_S and sulfane sulfur levels (***Supplemental Figure 6A and 6B***). While Survodutide effectively improved metabolic parameters, including reduced body weight, plasma and hepatic triglycerides, and plasma total cholesterol, it unexpectedly impaired glucose tolerance. This likely stems from the glucagon receptor agonism, which may not be completely counteracted by GLP-1 receptor stimulation. Importantly, the addition of DATS synergistically associated with improvements in metabolic parameters (***Figure 7B-7G***).

**Figure 7.**
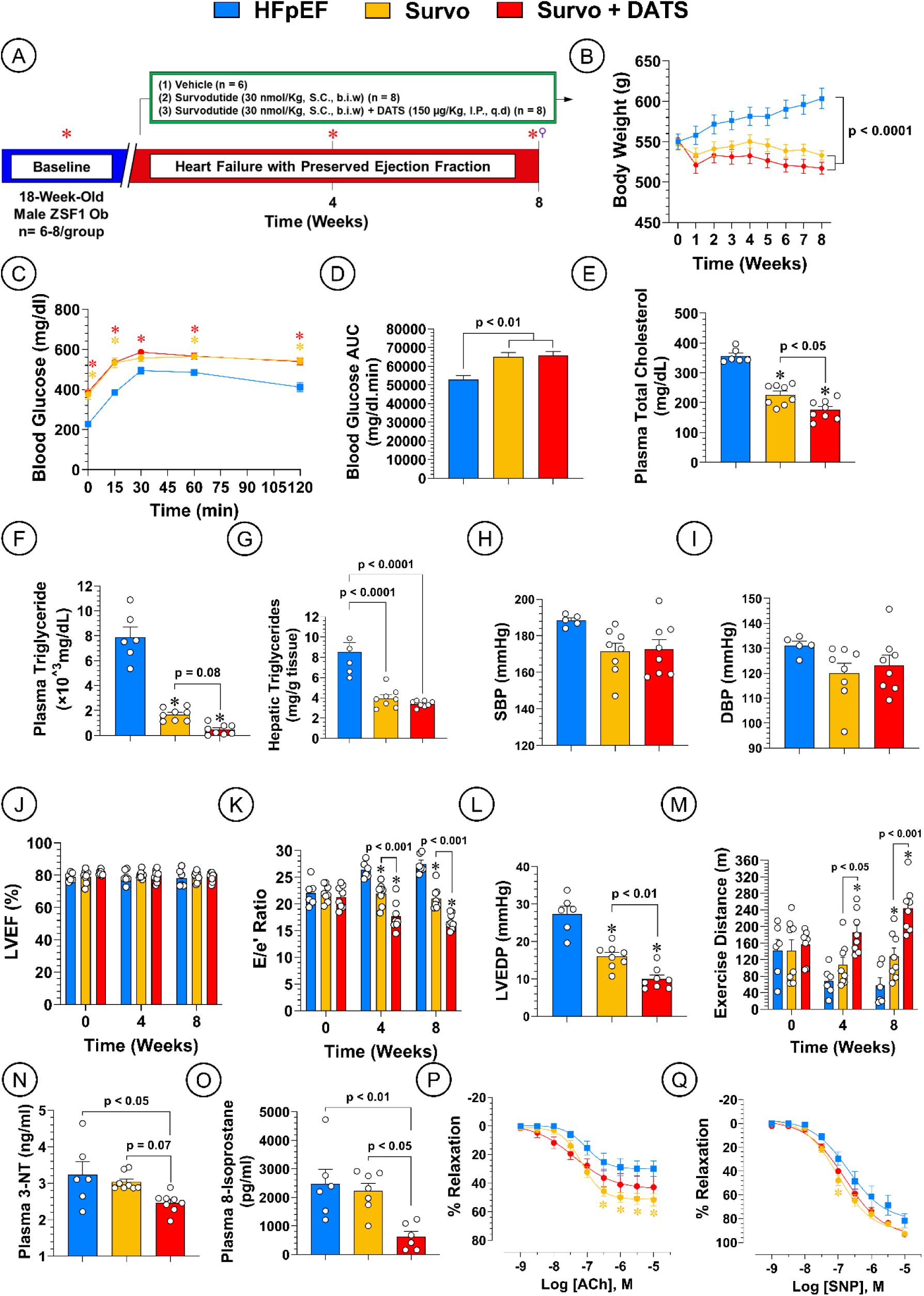
H_2_S Donor Synergizes with a dual GLP-1/glucagon agonist and Ameliorates HFpEF in the ZSF1 Obese Rat. ***(A)*** Study Timeline, (***B***) Body weight in grams, *(**C**)* Glucose tolerance test, GTT, *(**D**)* GTT area under the curve, *(**E**)* Plasma triglycerides, *(**F**)* Hepatic triglycerides, *(**G**)* Hepatic cholesterol, (***H***) Systolic blood pressure at 26^th^ week, *(**I**)* Diastolic blood pressure, *(**J**)* Left ventricular ejection fraction (LVEF), *(**K**)* Ratio of early mitral diastolic inflow velocity (E) and mitral annular early diastolic velocity (e’), (***L***) Left ventricular end diastolic pressure (LVEDP), *(**M**)* Treadmill exercise distance, (***N***) Plasma 3-nitrotrosine, *(**O**)* Plasma 8-isoprostane, *(**P**)* Aortic vascular reactivity to acetylcholine (ACh), ***(Q)*** Aortic vascular reactivity to sodium nitroprusside (SNP). Data in panels **C-I, L and N-Q** were collected at the last timepoint (week 8). Data in panels ***B, C, J, K, P and Q*** were analyzed with 2-way ANOVA followed by Sidak’s test. Data in panels ***D-I, L, N and O*** were analyzed with one-way ANOVA followed by Tukey test. Data are presented as mean ± SEM. * p < 0.05 vs. Control. * Echocardiography and Treadmill exercise, ⚲ Aortic vascular reactivity and Invasive hemodynamics.

Both Survodutide and combination therapy with DATS failed to significantly reduce systemic blood pressure or LVEF (***Figure 7H-7J***). Notably, Survodutide significantly improved diastolic function, as evidenced by reduced E/e’ and LVEDP, and enhanced exercise capacity. These beneficial effects were further augmented by DATS, resulting in a synergistic improvement in these parameters (***Figure 7K-7O***). Furthermore, both treatments significantly improved endothelium-dependent and -independent aortic vascular relaxation (**Figure 7P and 7Q**).

Remarkably, the combination therapy significantly reduced hepatic lipid content and attenuated cardiac interstitial and perivascular fibrosis and was superior to Survodutide treatment alone (***Figure 8 and Supplemental Figure 6C***). These findings highlight the therapeutic potential of H_2_S supplementation in HFpEF and demonstrate its ability to potentiate the effects of clinically relevant therapies like Survodutide.

**Figure 8.**
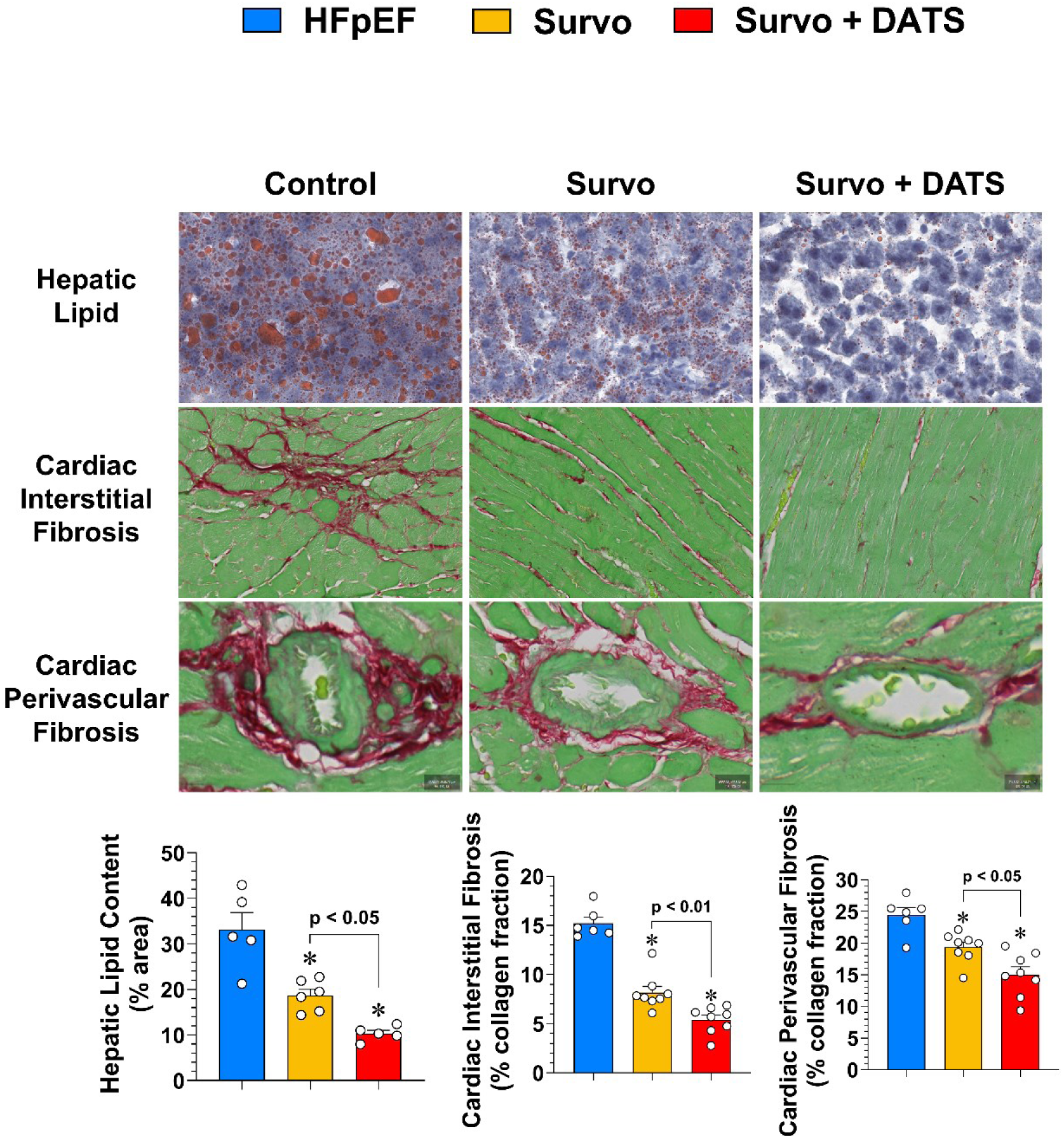
Survodutide and DATS Attenuate Hepatic Lipid Content and Cardiac Fibrosis in ZSF1 Obese Rat HFpEF Model. Representative images of hepatic lipid accumulation stained with Oil Red O, and representative images of cardiac fibrosis stained with Masson’s Trichrome and fast green counterstaining and respective quantification. Hepatic lipid content expressed as % area in bar graph. Cardiac interstitial and perivascular fibrosis expressed as % collagen fraction in bar graphs. Data were analyzed with one-way ANOVA followed by Tukey test. Data are presented as mean ± SEM.

## DISCUSSION

In this study, we identified systemic reductions in H_2_S as a potential contributor to the development of cardiometabolic HFpEF, evident in both human patients and two distinct preclinical models. Our data indicate that H_2_S deficiency and HFpEF pathology was remediated following the administration of pharmacological agents that release H_2_S. Our data show that H_2_S deficiency in HFpEF emerges from reduced tissue CSE expression and activity, and increased tissue SQR expression. In addition, we show that H_2_S synergizes with the dual GLP-1/glucagon RA, survodutide, in ameliorating HFpEF.

Our findings reveal a remarkable ∼80% reduction in circulating H_2_S levels in plasma from HFpEF patients compared to non-failing controls. This dramatic decrease in H_2_S bioavailability far exceeds the ∼40-50% reduction reported in HFrEF cases^15^, suggesting that H_2_S deficiency may play an equal or more significant role in HFpEF pathophysiology. Correspondingly, our preclinical models demonstrated a 60% reduction in circulating H_2_S in the two-hit mouse model and up to 90% in ZSF1 Ob rats. Notably, the ZSF1 Ob rat exhibits a more pronounced H_2_S deficiency that closely mirrors the clinical observations in HFpEF patients.

We further examined H_2_S levels across various organs including the heart, liver, and kidney given the systemic nature of HFpEF. In both the two-hit mouse model and ZSF1 Ob rats, we observed a decline in H_2_S levels in the plasma and peripheral orans. Utilizing the two-hit mouse model, we confirmed the presence of pronounced diastolic dysfunction, exercise intolerance, and metabolic dysregulation, consistent with previous studies^5,44^. The observed H_2_S deficiency was primarily attributable to decreased expression and activity of CSE in peripheral organs. While CSE is constitutively expressed in the liver, kidneys, vasculature, and heart, vascular endothelial cells as well as the liver and kidney^10,54^ are believed to be the primary regulators of circulating H_2_S levels. Our experiments with endothelial CSE knockout mice revealed that the depletion of this physiological sulfur pool exacerbated cardiac and vascular dysfunction. Conversely, exogenous H_2_S therapy with the H_2_S donor, JK-1, resulted in cardiovascular beneficial effects and marked improvements in exercise capacity. Indeed, previous studies show that CSE-derived H_2_S has multiple cardiovascular benefits including vasorelaxation^55-57^, angiogenesis regulation^56,58,59^, atheroprotection^60-63^ and reduced myocardial remodeling^22,43,64,65^. This study focused on CSE as an H_2_S producing enzyme as CBS is primarily localized to the central nervous system and has limited evidence of involvement in cardiovascular diseases^12^. The role of the mitochondrial 3-MST in HFpEF is currently under investigation by our group.

Our observations in ZSF1 Ob rats further corroborate the results from the two-hit model, as this model is characterized by a comprehensive phenotype that includes hypertension, obesity, diabetes, diastolic dysfunction, exercise intolerance, non-alcoholic fatty liver disease (NAFLD), and renal impairment^53,66,67^. Notably, the ZSF1 Ob rats exhibited even greater reductions in circulating, hepatic, and renal H_2_S levels, potentially elucidating the observed hepatic and renal pathologies in this model^66,67^. While both models share reductions in CSE expression, ZSF1 Ob rats demonstrated increased SQR expression. The later enzyme catalyzes the first irreversible step in the metabolism of H_2_S, and thus plays a pivotal role in the regulation of H_2_S-mediated signaling^12^. Previous data show that SQR inhibition attenuated hypertrophy, lung congestion, LV dilatation and myocardial fibrosis in HFrEF^68^. Future studies could explore the potential of SQR inhibitors as a therapeutic strategy for HFpEF.

While previous investigations have primarily focused on H_2_S effects within single organ systems, our findings underscore the importance of considering systemic dysregulated H_2_S signaling in HFpEF. The pleiotropic nature of H_2_S suggests that multiple mechanisms may contribute to the observed phenotype. Dysfunction in the liver and kidney, as observed in many HFpEF patients, can significantly impact overall H_2_S bioavailability. For example, hepatic dysfunction, often observed in patients with HFpEF, can lead to impaired H_2_S production and contribute to the systemic H_2_S deficiency observed in our study. This emphasizes the systemic nature of H_2_S dysregulation in HFpEF and highlights the importance of considering the contributions of multiple organs, beyond the heart, to the overall pathophysiology of this complex disease.

A key clinically relevant finding from our study is the synergistic benefits of H_2_S when combined with the dual GLP-1/glucagon RA, survodutide, given the complexity of cardiometabolic HFpEF that necessitates a multifaceted treatment approach. While current therapies such as SGLT2 inhibitors and mineralocorticoid receptor antagonists offer some benefit, they have limited impact on mortality in HFpEF patients^69,70^. This unmet need drives the search for novel therapies, including GLP-1RAs. Notably, this drug class is clinically used to treat the major comorbidities of HFpEF i.e., obesity and diabetes and recently have gained a considerable attention due to their potential efficacy in a plethora of many diseases. Emerging clinical investigations include cardiovascular diseases, metabolic liver disease, chronic kidney disease, sleep apnea, neurodegenerative disorders, addiction and obesity-associated cancers^71-73^.

The single GLP-1RA, semaglutide, significantly improved heart failure symptoms and physical limitations, and lowered the risk of major adverse outcomes of heart failure in obese HFpEF patients without (STEP-HFpEF trial) or with (STEP-HFpEF DM trial) type 2 diabetes^33,74^. Interestingly, semaglutide attenuated CRP levels and was associated with 10.7% and 6.4% reductions in body weight in both trials, respectively. Mechanistically, the beneficial effects of semaglutide in HFpEF was attributed to the extent of weight loss^75^, reducing inflammation^76^ and improving adverse cardiac remodeling^77^. Remarkably, a pooled analysis of four major semaglutide trials (SELECT, FLOW, STEP-HFpEF, and STEP-HFpEF DM) confirmed the substantial efficacy of this drug in heart failure^78^.

The remarkable findings of semaglutide in HFpEF encouraged the investigation of the dual GLP-1/GIP RA, tirzepatide, in this condition (SUMMIT trial)^32^. Similar to semaglutide, tirzepatide reduced CRP and body weight (11.6%), and was associated with reduced worsening heart-failure events and improved exercise tolerance. In addition to body weight reduction, other mechanisms for tirzepatide in HFpEF have been suggested, including reduced circulatory volume–pressure overload and systemic inflammation and mitigated cardiovascular–kidney end-organ injury^79^, and reducing paracardiac adipose tissue^80^.

The dual GLP-1/glucagon RA including survodutide, is a new emerging class under development and has not been explored in HFpEF. This class was discovered based on oxyntomodulin, an endogenous peptide hormone with a weak dual agonism on GLP-1 and glucagon receptors^81^, to simultaneously target both energy expenditure and food intake and induce weight loss. Survodutide has been examined in a phase 2 randomized, double-blind trial in persons with pre-obesity and obesity without diabetes and led to a dose-dependent reduction in body weight^82,83^. A post hoc analysis revealed that survodutide also improved both systolic and diastolic blood pressure^84^. In a 16-week trial, survodutide demonstrated superior efficacy to semaglutide in reducing HbA1c levels and body weight in patients with type 2 diabetes^85^. Recently, survodutide has been revealed to improve metabolic dysfunction-associated steatohepatitis (MASH) with significant improvements in fibrosis in a phase 2 randomized trial^34^. This drug is currently investigated in a phase 3, randomized trial (SYNCHRONIZE-CVOT) to determine its cardiovascular safety and efficacy^86^. Our study represents the first investigation into the beneficial effects of this class in HFpEF. Furthermore, while these drugs offer substantial potential, we hypothesize that combining them with H_2_S donors could synergistically enhance their therapeutic impact and provide a more comprehensive approach to combating this complex disease.

Our study indicates that H_2_S may further enhance the weight loss and metabolic benefits of GLP-1RAs. This is supported by observed improvements in lipid profiles and reduced hepatic lipid content. While previous research demonstrated that survodutide reduces both circulating and hepatic lipids in diet-induced obese mice, those experiments showed maintained glycemic control, a finding that differs from our results in ZSF1 Ob rats^87^. This discrepancy may be attributed to species-specific differences. ZSF1 Ob rats have higher baseline glucose levels compared to mice and could be more susceptible to the hyperglycemic effects of glucagon. Additionally, the previous study utilized a shorter subchronic design, unlike our study. Although clinical studies on survodutide have not specifically addressed glucose tolerance, they have shown a reduction in HbA1c levels^85^. The synergy of H_2_S with survodutide on improving metabolic parameters emerges from H_2_S pleiotropic actions in multiple organs and its effect on hepatic and lipid metabolism^88,89^.

This study reveals a striking synergy between survodutide and H_2_S in enhancing cardiac function, exercise tolerance, and reducing cardiac fibrosis in a ZSF1 Ob model of HFpEF. This synergistic effect is particularly noteworthy when compared to our previous investigation, which demonstrated a potentiation effect with the combination of the SGLT2 inhibitor empagliflozin and H_2_S in the same model. Several key findings emerge from this study. Firstly, survodutide alone demonstrated superior efficacy compared to empagliflozin, which had no significant impact on multiple parameters. Secondly, the combination of H_2_S and survodutide markedly improved exercise capacity, a clinically relevant marker. This combination led to progressive improvement in exercise capacity over time, whereas H_2_S plus empagliflozin only prevented deterioration without further enhancement. Crucially, this study administered survodutide and H_2_S during the advanced, severe stage of HFpEF (between 18 and 26 weeks of age). This contrasts with our previous study, where empagliflozin and H_2_S treatment was initiated earlier (10 to 18 weeks of age) before significant disease progression.

## STUDY LIMITATIONS

This study utilized male ZSF1 Ob rats and murine models of hypertension and obesity-induced HFpEF. While these rodent models are well-established and characterized, they primarily reflect metabolic dysfunction and blood pressure dysregulation, representing only a subset of HFpEF patients. Additionally, the exclusive use of male rats and mice limits the generalizability of our findings. Future research should focus on investigating changes in cardiac and systemic H_2_S bioavailability, as well as the effects of H_2_S therapy, in female animals across various HFpEF subtypes.

Another limitation is the use of only endothelial-specific CSE KO and Tg mice, while CSE is highly expressed in other organs like the liver and the kidney. Future studies should focus on global or organ specific CSE knockout.

A further limitation is that H_2_S donor therapy was administered early in the development and progression of HFpEF. Specifically, in the murine model, the H_2_S donor JK-1 was given five weeks after the initiation of L-NAME and a high-fat diet, a period during which H_2_S levels remain relatively normal. To rigorously evaluate H_2_S therapy’s efficacy in HFpEF, future experiments should involve administering H_2_S donors at later stages of HFpEF progression.

## CONCLUSIONS

This study offers novel insights into the dysregulation of H_2_S in cardiometabolic HFpEF. We observed significant reductions in H_2_S and sulfane sulfur levels in circulation, as well as in myocardial, hepatic, and renal tissues across multiple clinically relevant HFpEF models. These decreases are primarily attributed to diminished CSE expression and activity and increased SQR expression. Furthermore, enhancing H_2_S bioavailability through pharmacological administration markedly improved the HFpEF phenotype.

Future research should focus on the organ-specific effects of H_2_S modulation through genetic models targeting CSE, 3-MST, and SQR, as well as the precise mechanisms underlying the global and local benefits of H_2_S donation in HFpEF. Additionally, the efficacy of SQR inhibitors in HFpEF should be explored.

Our findings demonstrate that H_2_S donor therapy synergistically enhances the beneficial effects of GLP-1/glucagon RAs, such as Survodutide, in improving metabolic parameters, diastolic function, exercise capacity, and reducing oxidative stress and fibrosis in a preclinical model of HFpEF. This synergistic interaction suggests that combining H_2_S donors with GLP-1/glucagon RAa may represent a promising therapeutic strategy for the treatment of cardiometabolic HFpEF.

Given that cardiometabolic HFpEF is characterized by multi-organ dysfunction, there is an urgent need for innovative therapeutic strategies and a deeper understanding of its pathophysiological drivers. Our findings strongly support a central role for H_2_S deficiency in the pathology of cardiometabolic HFpEF, highlighting the necessity for further investigation into the role of H_2_S in this condition.

### Clinical Perspectives

#### Clinical competencies

This study highlights the critical role of hydrogen sulfide (H_2_S) in heart failure with preserved ejection fraction (HFpEF), underscoring several key competencies for clinicians. First, healthcare providers should recognize the potential for H_2_S deficiency in HFpEF patients, as this deficiency may contribute to disease progression and poor outcomes. Understanding the underlying pathophysiology—specifically the decreased production of H_2_S by cystathionine-γ-lyase (CSE) and its increased metabolism by sulfide quinone oxidoreductase (SQR)—can inform treatment strategies. Additionally, the findings suggest that pharmacological H_2_S supplementation may improve diastolic function and reduce cardiac fibrosis, encouraging clinicians to consider emerging therapies that target H_2_S bioavailability. Importantly, the study demonstrates a synergistic interaction between H_2_S supplementation and GLP-1/glucagon RAs in improving key HFpEF parameters. This finding has significant clinical implications, as it suggests that combining these two therapeutic approaches may offer a more effective treatment strategy than either agent alone for patients with HFpEF. This study ultimately emphasizes the importance of a holistic approach to managing HFpEF, integrating metabolic and inflammatory factors alongside traditional cardiac care.

#### Translational Outlook

While the study offers promising insights, several barriers must be addressed for successful clinical translation. Further research is needed to establish optimal dosing, delivery methods, and long-term safety of H_2_S donors in HFpEF patients. It is also crucial to explore how individual patient characteristics, including comorbidities and genetic factors, influence H_2_S metabolism and response to therapy. Robust clinical trials are essential to validate the efficacy of H_2_S supplementation across diverse populations with HFpEF, ensuring that findings are generalizable. Lastly, the integration of H_2_S-related interventions into existing clinical guidelines for HFpEF management will require collaboration among researchers, clinicians, and guideline committees. By addressing these barriers, future research can enhance clinical practice and improve outcomes for patients with HFpEF.

## ABBREVIATIONS AND ACRONYMS

CSE: cystathionine-γ-lyase
GLP-1: glucagon-like peptide 1
GLP-1RA: GLP-1 receptor agonist
HFpEF: heart failure with preserved ejection fraction
HFrEF: heart failure with reduced ejection fraction
L-NAME: N(ω)-nitro-L-arginine methyl ester
LV: left ventricle
SNP: sodium nitroprusside
SQR: sulfide quinone oxidoreductase
WKY: Wistar Kyoto
ZSF1: Zucker fatty and spontaneously hypertensive

## Acknowledgements

We thank Alexandra Nevins, Silpa Arkat, and Cell Biology and Bioimaging Core at Pennington Biomedical Research Center for technical support.

## Data Sharing Statement

The data underlying this article will be shared on reasonable request to the corresponding author.

## Notes

**Author/Funding disclosures:** These studies were supported by grants from the National Institutes of Health HL146098, HL146514, and HL151398 to D.J.L., HL159428 to T.T.G., AA029984 to T.E.S., AHA Postdoctoral Fellowship to Z.L. (20POST35200075), P20GM135002 and U54GM104940 to T.D.A., and NIH CCTS-Training Grant TL1TR003106 to J.D. D.J.L. serves as an unpaid consultant for Sulfagenix, Inc. The other authors declare no conflicts of interest.

### Competing Interest Statement

D.J.L. serves as an unpaid consultant for Sulfagenix, Inc. Sulfagenix is currently developing H2S-based therapeutics for a number of clinical indications including cardiovascular diseases. The other authors declare no conflicts of interest.

### Summary of Updates

New results including a combination therapy of an H2S donor with the GLP-1/glucagon receptor agonist, survodutide, in HFpEF.

## REFERENCES

1. Savarese G, Becher PM, Lund LH, Seferovic P, Rosano GMC, Coats AJS. Global burden of heart failure: a comprehensive and updated review of epidemiology. Cardiovasc Res. 2023;118:3272–3287. doi: 10.1093/cvr/cvac013

2. Loop MS, Van Dyke MK, Chen L, Brown TM, Durant RW, Safford MM, Levitan EB. Comparison of Length of Stay, 30-Day Mortality, and 30-Day Readmission Rates in Medicare Patients With Heart Failure and With Reduced Versus Preserved Ejection Fraction. Am J Cardiol. 2016;118:79-85. doi: 10.1016/j.amjcard.2016.04.015

3. Pfeffer MA, Shah AM, Borlaug BA. Heart Failure With Preserved Ejection Fraction In Perspective. Circulation Research. 2019;124:1598–1617. doi: 10.1161/circresaha.119.313572

4. Shah SJ, Borlaug BA, Kitzman DW, McCulloch AD, Blaxall BC, Agarwal R, Chirinos JA, Collins S, Deo RC, Gladwin MT, et al. Research Priorities for Heart Failure With Preserved Ejection Fraction. Circulation. 2020;141:1001–1026. doi: 10.1161/circulationaha.119.041886

5. Schiattarella GG, Altamirano F, Tong D, French KM, Villalobos E, Kim SY, Luo X, Jiang N, May HI, Wang ZV, et al. Nitrosative stress drives heart failure with preserved ejection fraction. Nature. 2019;568:351–356. doi: 10.1038/s41586-019-1100-z

6. Cruz L, Ryan JJ. Nitric Oxide Signaling in Heart Failure With Preserved Ejection Fraction. JACC Basic Transl Sci. 2017;2:341–343. doi: 10.1016/j.jacbts.2017.05.004

7. Pan LL, Qin M, Liu XH, Zhu YZ. The Role of Hydrogen Sulfide on Cardiovascular Homeostasis: An Overview with Update on Immunomodulation. Front Pharmacol. 2017;8:686. doi: 10.3389/fphar.2017.00686

8. Kolluru GK, Shackelford RE, Shen X, Dominic P, Kevil CG. Sulfide regulation of cardiovascular function in health and disease. Nat Rev Cardiol. 2023;20:109–125. doi: 10.1038/s41569-022-00741-6

9. Zhao W, Zhang J, Lu Y, Wang R. The vasorelaxant effect of H(2)S as a novel endogenous gaseous K(ATP) channel opener. EMBO J. 2001;20:6008–6016. doi: 10.1093/emboj/20.21.6008

10. Yang G, Wu L, Jiang B, Yang W, Qi J, Cao K, Meng Q, Mustafa AK, Mu W, Zhang S, et al. H2S as a Physiologic Vasorelaxant: Hypertension in Mice with Deletion of Cystathionine γ-Lyase. Science. 2008;322:587–590. doi: 10.1126/science.1162667

11. Peleli M, Bibli SI, Li Z, Chatzianastasiou A, Varela A, Katsouda A, Zukunft S, Bucci M, Vellecco V, Davos CH, et al. Cardiovascular phenotype of mice lacking 3-mercaptopyruvate sulfurtransferase. Biochem Pharmacol. 2020;176:113833. doi: 10.1016/j.bcp.2020.113833

12. Cirino G, Szabo C, Papapetropoulos A. Physiological roles of hydrogen sulfide in mammalian cells, tissues, and organs. Physiol Rev. 2023;103:31–276. doi: 10.1152/physrev.00028.2021

13. Lapenna KB, Polhemus DJ, Doiron JE, Hidalgo HA, Li Z, Lefer DJ. Hydrogen Sulfide as a Potential Therapy for Heart Failure—Past, Present, and Future. Antioxidants. 2021;10:485. doi: 10.3390/antiox10030485

14. Wang R. Roles of Hydrogen Sulfide in Hypertension Development and Its Complications: What, So What, Now What. Hypertension. 2023;80:936–944. doi: 10.1161/hypertensionaha.122.19456

15. Polhemus DJ, Calvert JW, Butler J, Lefer DJ. The Cardioprotective Actions of Hydrogen Sulfide in Acute Myocardial Infarction and Heart Failure. Scientifica. 2014;2014:1–8. doi: 10.1155/2014/768607

16. Polhemus DJ, Lefer DJ. Emergence of Hydrogen Sulfide as an Endogenous Gaseous Signaling Molecule in Cardiovascular Disease. Circulation Research. 2014;114:730–737. doi: 10.1161/circresaha.114.300505

17. Elrod JW, Calvert JW, Morrison J, Doeller JE, Kraus DW, Tao L, Jiao X, Scalia R, Kiss L, Szabo C, et al. Hydrogen sulfide attenuates myocardial ischemia-reperfusion injury by preservation of mitochondrial function. Proceedings of the National Academy of Sciences. 2007;104:15560–15565. doi: 10.1073/pnas.0705891104

18. Calvert JW, Jha S, Gundewar S, Elrod JW, Ramachandran A, Pattillo CB, Kevil CG, Lefer DJ. Hydrogen sulfide mediates cardioprotection through Nrf2 signaling. Circ Res. 2009;105:365–374. doi: 10.1161/CIRCRESAHA.109.199919

19. Li Z, Xia H, Sharp TE, Lapenna KB, Elrod JW, Casin KM, Liu K, Calvert JW, Chau VQ, Salloum FN, et al. Mitochondrial H2S Regulates BCAA Catabolism in Heart Failure. Circulation Research. 2022;131:222–235. doi: 10.1161/circresaha.121.319817

20. Zanardo RCO, Brancaleone V, Distrutti E, Fiorucci S, Cirino G, Wallace JL. Hydrogen Sulfide is an Endogenous Modulator of Leukocyte-mediated Inflammation. The FASEB Journal. 2006;20:2118–2120. doi: 10.1096/fj.06-6270fje

21. Mustafa AK, Gadalla MM, Sen N, Kim S, Mu W, Gazi SK, Barrow RK, Yang G, Wang R, Snyder SH. H2S Signals Through Protein S-Sulfhydration. Science Signaling. 2009;2:ra72-ra72. doi: 10.1126/scisignal.2000464

22. King AL, Polhemus DJ, Bhushan S, Otsuka H, Kondo K, Nicholson CK, Bradley JM, Islam KN, Calvert JW, Tao Y-X, et al. Hydrogen sulfide cytoprotective signaling is endothelial nitric oxide synthase-nitric oxide dependent. Proceedings of the National Academy of Sciences. 2014;111:3182–3187. doi: 10.1073/pnas.1321871111

23. Capone F, Vettor R, Schiattarella GG. Cardiometabolic HFpEF: NASH of the Heart. Circulation. 2023;147:451–453. doi: doi:10.1161/CIRCULATIONAHA.122.062874

24. Shah SJ, Katz DH, Selvaraj S, Burke MA, Yancy CW, Gheorghiade M, Bonow RO, Huang C-C, Deo RC. Phenomapping for Novel Classification of Heart Failure With Preserved Ejection Fraction. Circulation. 2015;131:269–279. doi: 10.1161/circulationaha.114.010637

25. Cohen JB, Schrauben SJ, Zhao L, Basso MD, Cvijic ME, Li Z, Yarde M, Wang Z, Bhattacharya PT, Chirinos DA, et al. Clinical Phenogroups in Heart Failure With Preserved Ejection Fraction. JACC: Heart Failure. 2020;8:172–184. doi: 10.1016/j.jchf.2019.09.009

26. Xu W, Cui C, Cui C, Chen Z, Zhang H, Cui Q, Xu G, Fan J, Han Y, Tang L, et al. Hepatocellular cystathionine gamma lyase/hydrogen sulfide attenuates nonalcoholic fatty liver disease by activating farnesoid X receptor. Hepatology. 2022;76:1794–1810. doi: 10.1002/hep.32577

27. Barr LA, Shimizu Y, Lambert JP, Nicholson CK, Calvert JW. Hydrogen sulfide attenuates high fat diet-induced cardiac dysfunction via the suppression of endoplasmic reticulum stress. Nitric Oxide. 2015;46:145–156. doi: 10.1016/j.niox.2014.12.013

28. Suzuki K, Olah G, Modis K, Coletta C, Kulp G, Gerö D, Szoleczky P, Chang T, Zhou Z, Wu L, et al. Hydrogen sulfide replacement therapy protects the vascular endothelium in hyperglycemia by preserving mitochondrial function. Proceedings of the National Academy of Sciences. 2011;108:13829–13834. doi: 10.1073/pnas.1105121108

29. Padiya R, Khatua TN, Bagul PK, Kuncha M, Banerjee SK. Garlic improves insulin sensitivity and associated metabolic syndromes in fructose fed rats. Nutrition & Metabolism. 2011;8:53. doi: 10.1186/1743-7075-8-53

30. Zhong G, Chen F, Cheng Y, Tang C, Du J. The role of hydrogen sulfide generation in the pathogenesis of hypertension in rats induced by inhibition of nitric oxide synthase. Journal of Hypertension. 2003;21:1879–1885.

31. Solini A, Trico D, Del Prato S. Incretins and cardiovascular disease: to the heart of type 2 diabetes? Diabetologia. 2023;66:1820–1831. doi: 10.1007/s00125-023-05973-w

32. Packer M, Zile MR, Kramer CM, Baum SJ, Litwin SE, Menon V, Ge J, Weerakkody GJ, Ou Y, Bunck MC, et al. Tirzepatide for Heart Failure with Preserved Ejection Fraction and Obesity. N Engl J Med. 2024. doi: 10.1056/NEJMoa2410027

33. Kosiborod MN, Abildstrom SZ, Borlaug BA, Butler J, Rasmussen S, Davies M, Hovingh GK, Kitzman DW, Lindegaard ML, Moller DV, et al. Semaglutide in Patients with Heart Failure with Preserved Ejection Fraction and Obesity. N Engl J Med. 2023;389:1069–1084. doi: 10.1056/NEJMoa2306963

34. Sanyal AJ, Bedossa P, Fraessdorf M, Neff GW, Lawitz E, Bugianesi E, Anstee QM, Hussain SA, Newsome PN, Ratziu V, et al. A Phase 2 Randomized Trial of Survodutide in MASH and Fibrosis. N Engl J Med. 2024;391:311–319. doi: 10.1056/NEJMoa2401755

35. Jastreboff AM, Kaplan LM, Frias JP, Wu Q, Du Y, Gurbuz S, Coskun T, Haupt A, Milicevic Z, Hartman ML, Retatrutide Phase 2 Obesity Trial I. Triple-Hormone-Receptor Agonist Retatrutide for Obesity - A Phase 2 Trial. N Engl J Med. 2023;389:514-526. doi: 10.1056/NEJMoa2301972

36. Urva S, Coskun T, Loh MT, Du Y, Thomas MK, Gurbuz S, Haupt A, Benson CT, Hernandez-Illas M, D’Alessio DA, Milicevic Z. LY3437943, a novel triple GIP, GLP-1, and glucagon receptor agonist in people with type 2 diabetes: a phase 1b, multicentre, double-blind, placebo-controlled, randomised, multiple-ascending dose trial. Lancet. 2022;400:1869-1881. doi: 10.1016/S0140-6736(22)02033-5

37. Li Z, Xia H, Sharp TE, Lapenna KB, Katsouda A, Elrod JW, Pfeilschifter J, Beck K-F, Xu S, Xian M, et al. Hydrogen Sulfide Modulates Endothelial–Mesenchymal Transition in Heart Failure. Circulation Research. 2023;132:154–166. doi: 10.1161/circresaha.122.321326

38. Xia H, Li Z, Sharp TE, Polhemus DJ, Carnal J, Moles KH, Tao YX, Elrod J, Pfeilschifter J, Beck KF, Lefer DJ. Endothelial Cell Cystathionine γ-Lyase Expression Level Modulates Exercise Capacity, Vascular Function, and Myocardial Ischemia Reperfusion Injury. Journal of the American Heart Association. 2020;9. doi: 10.1161/jaha.120.017544

39. Kang J, Li Z, Organ CL, Park CM, Yang CT, Pacheco A, Wang D, Lefer DJ, Xian M. pH-Controlled Hydrogen Sulfide Release for Myocardial Ischemia-Reperfusion Injury. J Am Chem Soc. 2016;138:6336–6339. doi: 10.1021/jacs.6b01373

40. Li Z, Organ CL, Kang J, Polhemus DJ, Trivedi RK, Sharp TE, 3rd, Jenkins JS, Tao YX, Xian M, Lefer DJ. Hydrogen Sulfide Attenuates Renin Angiotensin and Aldosterone Pathological Signaling to Preserve Kidney Function and Improve Exercise Tolerance in Heart Failure. JACC Basic Transl Sci. 2018;3:796–809. doi: 10.1016/j.jacbts.2018.08.011

41. Li Z, Xia H, Sharp TE, 3rd, LaPenna KB, Elrod JW, Casin KM, Liu K, Calvert JW, Chau VQ, Salloum FN, et al. Mitochondrial H(2)S Regulates BCAA Catabolism in Heart Failure. Circ Res. 2022;131:222-235. doi: 10.1161/CIRCRESAHA.121.319817

42. Li Z, Xia H, Sharp TE, 3rd, LaPenna KB, Katsouda A, Elrod JW, Pfeilschifter J, Beck KF, Xu S, Xian M, et al. Hydrogen Sulfide Modulates Endothelial-Mesenchymal Transition in Heart Failure. Circ Res. 2023;132:154–166. doi: 10.1161/CIRCRESAHA.122.321326

43. Polhemus D, Kondo K, Bhushan S, Bir SC, Kevil CG, Murohara T, Lefer DJ, Calvert JW. Hydrogen sulfide attenuates cardiac dysfunction after heart failure via induction of angiogenesis. Circ Heart Fail. 2013;6:1077–1086. doi: 10.1161/CIRCHEARTFAILURE.113.000299

44. Lapenna KB, Li Z, Doiron JE, Sharp TE, Xia H, Moles K, Koul K, Wang JS, Polhemus DJ, Goodchild TT, et al. Combination Sodium Nitrite and Hydralazine Therapy Attenuates Heart Failure With Preserved Ejection Fraction Severity in a “2-Hit” Murine Model. Journal of the American Heart Association. 2023. doi: 10.1161/jaha.122.028480

45. Mok YY, Atan MS, Yoke Ping C, Zhong Jing W, Bhatia M, Moochhala S, Moore PK. Role of hydrogen sulphide in haemorrhagic shock in the rat: protective effect of inhibitors of hydrogen sulphide biosynthesis. Br J Pharmacol. 2004;143:881–889. doi: 10.1038/sj.bjp.0706014

46. Mehlem A, Hagberg CE, Muhl L, Eriksson U, Falkevall A. Imaging of neutral lipids by oil red O for analyzing the metabolic status in health and disease. Nat Protoc. 2013;8:1149–1154. doi: 10.1038/nprot.2013.055

47. Pugliese NR, Pellicori P, Filidei F, De Biase N, Maffia P, Guzik TJ, Masi S, Taddei S, Cleland JGF. Inflammatory pathways in heart failure with preserved left ventricular ejection fraction: implications for future interventions. Cardiovasc Res. 2023;118:3536–3555. doi: 10.1093/cvr/cvac133

48. Gemici B, Wallace JL. Anti-inflammatory and cytoprotective properties of hydrogen sulfide. Methods Enzymol. 2015;555:169–193. doi: 10.1016/bs.mie.2014.11.034

49. Cao X, Ding L, Xie ZZ, Yang Y, Whiteman M, Moore PK, Bian JS. A Review of Hydrogen Sulfide Synthesis, Metabolism, and Measurement: Is Modulation of Hydrogen Sulfide a Novel Therapeutic for Cancer? Antioxid Redox Signal. 2019;31:1–38. doi: 10.1089/ars.2017.7058

50. Nayor M, Houstis NE, Namasivayam M, Rouvina J, Hardin C, Shah RV, Ho JE, Malhotra R, Lewis GD. Impaired Exercise Tolerance in Heart Failure With Preserved Ejection Fraction: Quantification of Multiorgan System Reserve Capacity. JACC Heart Fail. 2020;8:605–617. doi: 10.1016/j.jchf.2020.03.008

51. Fudim M, Zhong L, Patel KV, Khera R, Abdelmalek MF, Diehl AM, McGarrah RW, Molinger J, Moylan CA, Rao VN, et al. Nonalcoholic Fatty Liver Disease and Risk of Heart Failure Among Medicare Beneficiaries. J Am Heart Assoc. 2021;10:e021654. doi: 10.1161/JAHA.121.021654

52. Roh J, Hill JA, Singh A, Valero-Muñoz M, Sam F. Heart Failure With Preserved Ejection Fraction: Heterogeneous Syndrome, Diverse Preclinical Models. Circulation Research. 2022;130:1906–1925. doi: 10.1161/circresaha.122.320257

53. Hamdani N, Franssen C, Lourenco A, Falcao-Pires I, Fontoura D, Leite S, Plettig L, Lopez B, Ottenheijm CA, Becher PM, et al. Myocardial titin hypophosphorylation importantly contributes to heart failure with preserved ejection fraction in a rat metabolic risk model. Circ Heart Fail. 2013;6:1239–1249. doi: 10.1161/CIRCHEARTFAILURE.113.000539

54. Cirino G, Szabo C, Papapetropoulos A. Physiological roles of hydrogen sulfide in mammalian cells, tissues, and organs. Physiological Reviews. 2023;103:31–276. doi: 10.1152/physrev.00028.2021

55. Cheng Y, Ndisang JF, Tang G, Cao K, Wang R. Hydrogen sulfide-induced relaxation of resistance mesenteric artery beds of rats. Am J Physiol Heart Circ Physiol. 2004;287:H2316–2323. doi: 10.1152/ajpheart.00331.2004

56. Coletta C, Papapetropoulos A, Erdelyi K, Olah G, Modis K, Panopoulos P, Asimakopoulou A, Gero D, Sharina I, Martin E, Szabo C. Hydrogen sulfide and nitric oxide are mutually dependent in the regulation of angiogenesis and endothelium-dependent vasorelaxation. Proc Natl Acad Sci U S A. 2012;109:9161–9166. doi: 10.1073/pnas.1202916109

57. Greaney JL, Kutz JL, Shank SW, Jandu S, Santhanam L, Alexander LM. Impaired Hydrogen Sulfide-Mediated Vasodilation Contributes to Microvascular Endothelial Dysfunction in Hypertensive Adults. Hypertension. 2017;69:902–909. doi: 10.1161/HYPERTENSIONAHA.116.08964

58. Altaany Z, Yang G, Wang R. Crosstalk between hydrogen sulfide and nitric oxide in endothelial cells. J Cell Mol Med. 2013;17:879–888. doi: 10.1111/jcmm.12077

59. Saha S, Chakraborty PK, Xiong X, Dwivedi SK, Mustafi SB, Leigh NR, Ramchandran R, Mukherjee P, Bhattacharya R. Cystathionine beta-synthase regulates endothelial function via protein S-sulfhydration. FASEB J. 2016;30:441–456. doi: 10.1096/fj.15-278648

60. Bibli SI, Hu J, Sigala F, Wittig I, Heidler J, Zukunft S, Tsilimigras DI, Randriamboavonjy V, Wittig J, Kojonazarov B, et al. Cystathionine gamma Lyase Sulfhydrates the RNA Binding Protein Human Antigen R to Preserve Endothelial Cell Function and Delay Atherogenesis. Circulation. 2019;139:101–114. doi: 10.1161/CIRCULATIONAHA.118.034757

61. Cheung SH, Kwok WK, To KF, Lau JY. Anti-atherogenic effect of hydrogen sulfide by over-expression of cystathionine gamma-lyase (CSE) gene. PLoS One. 2014;9:e113038. doi: 10.1371/journal.pone.0113038

62. Mani S, Li H, Untereiner A, Wu L, Yang G, Austin RC, Dickhout JG, Lhotak S, Meng QH, Wang R. Decreased endogenous production of hydrogen sulfide accelerates atherosclerosis. Circulation. 2013;127:2523–2534. doi: 10.1161/CIRCULATIONAHA.113.002208

63. Zhang H, Guo C, Wu D, Zhang A, Gu T, Wang L, Wang C. Hydrogen sulfide inhibits the development of atherosclerosis with suppressing CX3CR1 and CX3CL1 expression. PLoS One. 2012;7:e41147. doi: 10.1371/journal.pone.0041147

64. Calvert JW, Elston M, Nicholson CK, Gundewar S, Jha S, Elrod JW, Ramachandran A, Lefer DJ. Genetic and pharmacologic hydrogen sulfide therapy attenuates ischemia-induced heart failure in mice. Circulation. 2010;122:11–19. doi: 10.1161/CIRCULATIONAHA.109.920991

65. Kondo K, Bhushan S, King AL, Prabhu SD, Hamid T, Koenig S, Murohara T, Predmore BL, Gojon G, Sr., Gojon G, Jr., et al. H(2)S protects against pressure overload-induced heart failure via upregulation of endothelial nitric oxide synthase. Circulation. 2013;127:1116–1127. doi: 10.1161/CIRCULATIONAHA.112.000855

66. Stolina M, Luo X, Dwyer D, Han C-Y, Chen R, Zhang Y, Xiong Y, Chen Y, Yin J, Shkumatov A, et al. The evolving systemic biomarker milieu in obese ZSF1 rat model of human cardiometabolic syndrome: Characterization of the model and cardioprotective effect of GDF15. PLOS ONE. 2020;15:e0231234. doi: 10.1371/journal.pone.0231234

67. Borges Canha M, Portela-Cidade JP, Conceicao G, Sousa-Mendes C, Leite S, Fontoura D, Moreira-Goncalves D, Falcao-Pires I, Lourenco A, Leite-Moreira A, Pimentel-Nunes P. Characterization of liver changes in ZSF1 rats, an animal model of metabolic syndrome. Rev Esp Enferm Dig. 2017;109:491–497. doi: 10.17235/reed.2017.4575/2016

68. Jackson MR, Cox KD, Baugh SDP, Wakeen L, Rashad AA, Lam PYS, Polyak B, Jorns MS. Discovery of a first-in-class inhibitor of sulfide:quinone oxidoreductase that protects against adverse cardiac remodelling and heart failure. Cardiovasc Res. 2022;118:1771–1784. doi: 10.1093/cvr/cvab206

69. Kapelios CJ, Murrow JR, Nuhrenberg TG, Montoro Lopez MN. Effect of mineralocorticoid receptor antagonists on cardiac function in patients with heart failure and preserved ejection fraction: a systematic review and meta-analysis of randomized controlled trials. Heart Fail Rev. 2019;24:367–377. doi: 10.1007/s10741-018-9758-0

70. Jaiswal A, Jaiswal V, Ang SP, Hanif M, Vadhera A, Agrawal V, Kumar T, Nair AM, Borra V, Garimella V, et al. SGLT2 inhibitors among patients with heart failure with preserved ejection fraction: A meta-analysis of randomised controlled trials. Medicine (Baltimore). 2023;102:e34693. doi: 10.1097/MD.0000000000034693

71. Alfaris N, Waldrop S, Johnson V, Boaventura B, Kendrick K, Stanford FC. GLP-1 single, dual, and triple receptor agonists for treating type 2 diabetes and obesity: a narrative review. EClinicalMedicine. 2024;75:102782. doi: 10.1016/j.eclinm.2024.102782

72. Mariam Z, Niazi SK. Glucagon-like peptide agonists: A prospective review. Endocrinol Diabetes Metab. 2024;7:e462. doi: 10.1002/edm2.462

73. Zheng Z, Zong Y, Ma Y, Tian Y, Pang Y, Zhang C, Gao J. Glucagon-like peptide-1 receptor: mechanisms and advances in therapy. Signal Transduct Target Ther. 2024;9:234. doi: 10.1038/s41392-024-01931-z

74. Kosiborod MN, Petrie MC, Borlaug BA, Butler J, Davies MJ, Hovingh GK, Kitzman DW, Moller DV, Treppendahl MB, Verma S, et al. Semaglutide in Patients with Obesity-Related Heart Failure and Type 2 Diabetes. N Engl J Med. 2024;390:1394–1407. doi: 10.1056/NEJMoa2313917

75. Borlaug BA, Kitzman DW, Davies MJ, Rasmussen S, Barros E, Butler J, Einfeldt MN, Hovingh GK, Moller DV, Petrie MC, et al. Semaglutide in HFpEF across obesity class and by body weight reduction: a prespecified analysis of the STEP-HFpEF trial. Nat Med. 2023;29:2358–2365. doi: 10.1038/s41591-023-02526-x

76. Verma S, Petrie MC, Borlaug BA, Butler J, Davies MJ, Kitzman DW, Shah SJ, Ronnback C, Abildstrom SZ, Liisberg K, et al. Inflammation in Obesity-Related HFpEF: The STEP-HFpEF Program. J Am Coll Cardiol. 2024;84:1646–1662. doi: 10.1016/j.jacc.2024.08.028

77. Solomon SD, Ostrominski JW, Wang X, Shah SJ, Borlaug BA, Butler J, Davies MJ, Kitzman DW, Verma S, Abildstrom SZ, et al. Effect of Semaglutide on Cardiac Structure and Function in Patients With Obesity-Related Heart Failure. J Am Coll Cardiol. 2024;84:1587–1602. doi: 10.1016/j.jacc.2024.08.021

78. Kosiborod MN, Deanfield J, Pratley R, Borlaug BA, Butler J, Davies MJ, Emerson SS, Kahn SE, Kitzman DW, Lingvay I, et al. Semaglutide versus placebo in patients with heart failure and mildly reduced or preserved ejection fraction: a pooled analysis of the SELECT, FLOW, STEP-HFpEF, and STEP-HFpEF DM randomised trials. Lancet. 2024;404:949–961. doi: 10.1016/S0140-6736(24)01643-X

79. Borlaug BA, Zile MR, Kramer CM, Baum SJ, Hurt K, Litwin SE, Murakami M, Ou Y, Upadhyay N, Packer M. Effects of tirzepatide on circulatory overload and end-organ damage in heart failure with preserved ejection fraction and obesity: a secondary analysis of the SUMMIT trial. Nat Med. 2024. doi: 10.1038/s41591-024-03374-z

80. Kramer CM, Borlaug BA, Zile MM, Ruff D, DiMaria JM, Menon V, Ou Y, Zarante AM, Hurt KC, Murakami M, et al. Tirzepatide Reduces LV Mass and Paracardiac Adipose Tissue in Obesity-Related Heart Failure: SUMMIT CMR Substudy. J Am Coll Cardiol. 2024. doi: 10.1016/j.jacc.2024.11.001

81. Clemmensen C, Finan B, Muller TD, DiMarchi RD, Tschop MH, Hofmann SM. Emerging hormonal-based combination pharmacotherapies for the treatment of metabolic diseases. Nat Rev Endocrinol. 2019;15:90–104. doi: 10.1038/s41574-018-0118-x

82. le Roux CW, Steen O, Lucas KJ, Startseva E, Unseld A, Hennige AM. Glucagon and GLP-1 receptor dual agonist survodutide for obesity: a randomised, double-blind, placebo-controlled, dose-finding phase 2 trial. Lancet Diabetes Endocrinol. 2024;12:162–173. doi: 10.1016/S2213-8587(23)00356-X

83. le Roux CW, Steen O, Lucas KJ, Startseva E, Unseld A, Hussain SA, Hennige AM. Subgroup analysis by sex and baseline BMI in people with a BMI >/=27 kg/m(2) in the phase 2 trial of survodutide, a glucagon/GLP-1 receptor dual agonist. Diabetes Obes Metab. 2025. doi: 10.1111/dom.16167

84. le Roux CW, Steen O, Lucas KJ, Ekinci EI, Startseva E, Unseld A, Hussain SA, Hennige AM. Survodutide, a glucagon receptor/glucagon-like peptide-1 receptor dual agonist, improves blood pressure in adults with obesity: A post hoc analysis from a randomized, placebo-controlled, dose-finding, phase 2 trial. Diabetes Obes Metab. 2025;27:993–996. doi: 10.1111/dom.16052

85. Bluher M, Rosenstock J, Hoefler J, Manuel R, Hennige AM. Dose-response effects on HbA(1c) and bodyweight reduction of survodutide, a dual glucagon/GLP-1 receptor agonist, compared with placebo and open-label semaglutide in people with type 2 diabetes: a randomised clinical trial. Diabetologia. 2024;67:470–482. doi: 10.1007/s00125-023-06053-9

86. Kosiborod MN, Platz E, Wharton S, le Roux CW, Brueckmann M, Ajaz Hussain S, Unseld A, Startseva E, Kaplan LM, Committees S-CT, Investigators. Survodutide for the Treatment of Obesity: Rationale and Design of the SYNCHRONIZE Cardiovascular Outcomes Trial. JACC Heart Fail. 2024;12:2101–2109. doi: 10.1016/j.jchf.2024.09.004

87. Zimmermann T, Thomas L, Baader-Pagler T, Haebel P, Simon E, Reindl W, Bajrami B, Rist W, Uphues I, Drucker DJ, et al. BI 456906: Discovery and preclinical pharmacology of a novel GCGR/GLP-1R dual agonist with robust anti-obesity efficacy. Mol Metab. 2022;66:101633. doi: 10.1016/j.molmet.2022.101633

88. Flori L, Piragine E, Calderone V, Testai L. Role of hydrogen sulfide in the regulation of lipid metabolism: Implications on cardiovascular health. Life Sci. 2024;341:122491. doi: 10.1016/j.lfs.2024.122491

89. Loiselle JJ, Yang G, Wu L. Hydrogen sulfide and hepatic lipid metabolism - a critical pairing for liver health. Br J Pharmacol. 2020;177:757–768. doi: 10.1111/bph.14556

