## Supplemental Material for "Hydrogen Sulfide Deficiency and Therapeutic Targeting in Cardiometabolic HFpEF: Evidence for Synergistic Benefit with GLP-1/Glucagon Agonism"

**Supplemental Appendix:**


Supplemental Table 1. Real-Time PCR Primer sequences Page 2
Supplemental Figure 1. HFpEF Pathology in "Two-hit" Murine HFpEF Model Page 3
Supplemental Figure 2. Hepatic & Renal Hydrogen Sulfide in "Two-Hit" HFpEF Mice Page 5
Supplemental Figure 3. Cardiac Function and Survival in HFpEF Control and EC-CSE Tg Mice Page 7
Supplemental Figure 5. Hydrogen Sulfide Therapy Attenuates HFpEF Severity in Murine "Two-Hit" Model Page 8
Supplemental Figure 6. Fibrosis in the ZSF1 Obese Rat Tissues Page 10
Supplemental Figure 7. Circulating H2S and Hepatic Lipid Content in ZSF1 Obese Rats treated with Survodutide and DATS Page 12

**Supplemental Table 1. Real-Time PCR Primer sequences**

| Gene | | Primer Sequences (5′–3′) |
|---|---|---|
| Mouse 18s | sense | CTCTAGATAACCTCGGGCC |
| | antisense | GAACCCTGATTCCCCGTCA |
| Mouse Tubb5 | sense | GGAAATCGTGCACATCCAGG |
| | antisense | GGGGTCGATGCCATGTTCAT |
| Mouse CSE | sense | GAGCCTGAGCAATGGAAT |
| | antisense | GATGGGTAATCGTAATGGTG |
| Mouse SQR | sense | TCATCGCTCTTGGAATCCAGC |
| | antisense | CATTGCCTTCCTTGAAACCCTG |
| Rat 18s | sense | GTAAACCCGTTGAACCCCATT |
| | antisense | CCATCCAATCGGTAGTAGCG |
| Rat CSE | sense | CAATGGAGTTCGCGTGCTG |
| | antisense | TTTCCAAGCAATTCCTCGTC |
| Rat SQR | sense | ATGGCCACTCGCATGAAGAGG |
| | antisense | GATGAGGACAGCTCTTTGGCAC |

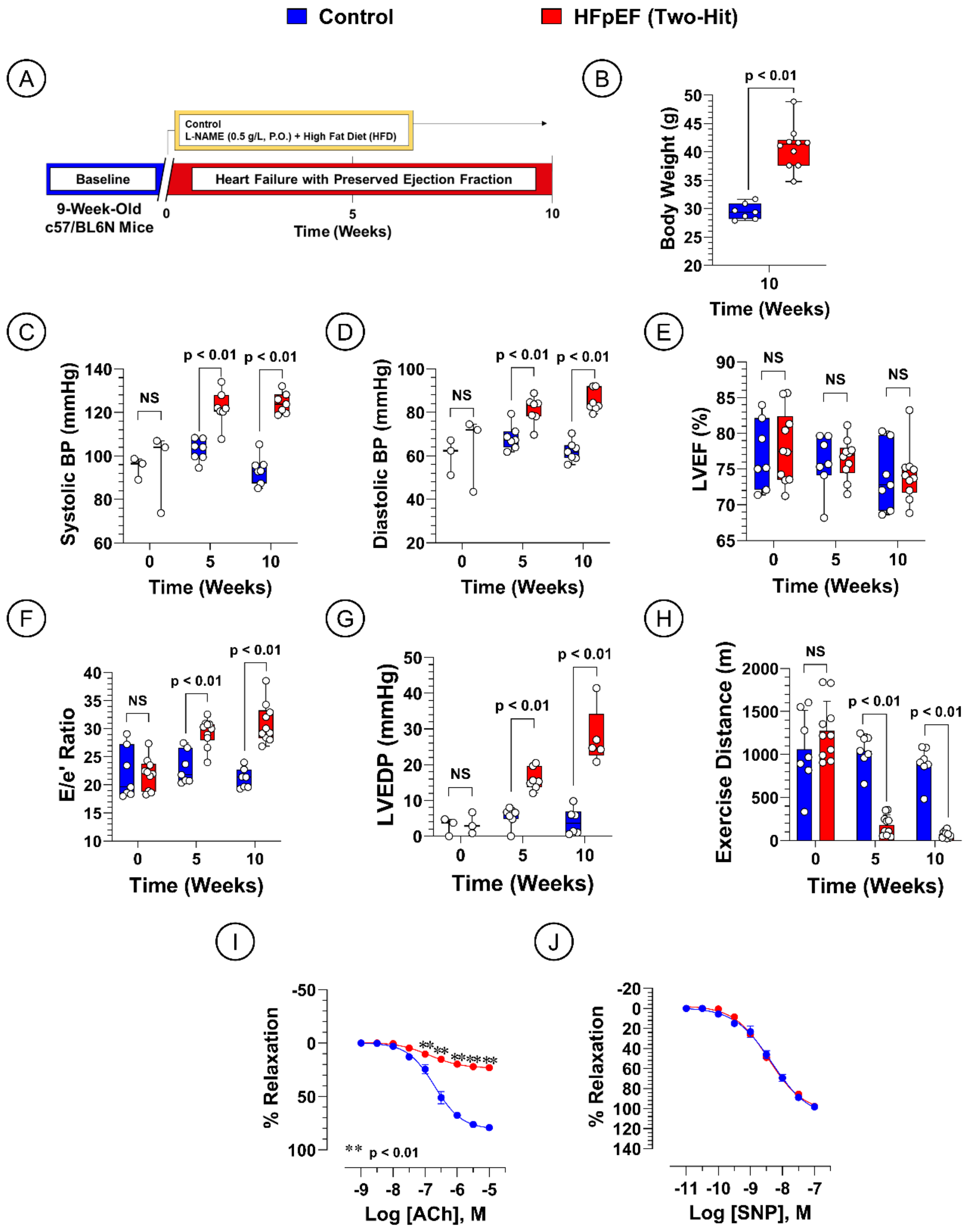


***Supplemental Figure 1. HFpEF Pathology in "Two-hit" Murine HFpEF Model***

(***A***) Study timeline, (***B***) Body weight, (***C***) Systolic blood pressure, (***D***) Diastolic blood pressure, ***(E)*** Left ventricular ejection fraction (LVEF), ***(F)*** Ratio of early mitral diastolic inflow velocity (E) and mitral annular early diastolic velocity (e'), (***G***) Left ventricular end diastolic pressure

(LVEDP), ***(H)*** Treadmill exercise distance, ***(I)*** Aortic vascular reactivity to acetylcholine (ACh) at week 10, ***(J)*** Aortic vascular reactivity to sodium nitroprusside (SNP) at week 10. Data in panel ***B*** was analyzed with Student unpaired 2-tail *t* test. Data in panels ***B-H*** were analyzed with multiple Student unpaired 2-tail *t* tests. Data in panels ***I*** and ***J*** were analyzed with ordinary 2-way ANOVA followed by Sidak's test. Data in panels ***B-G*** are presented as box plots (median, minimum, and maximum). Data in panels ***H-J*** are presented as mean ± SEM.

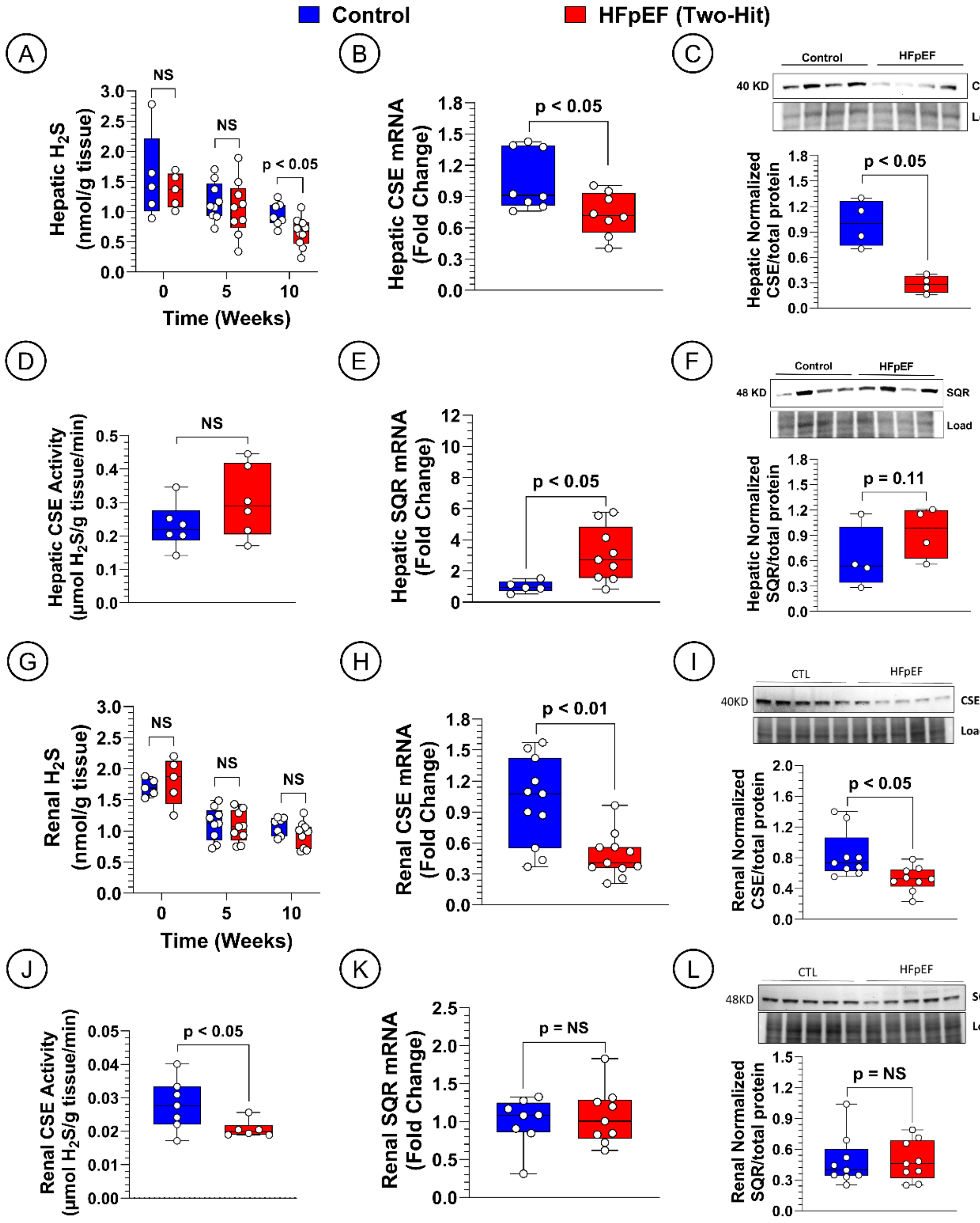


***Supplemental Figure 2. Hepatic and Renal Hydrogen Sulfide in "Two-Hit" HFpEF Mice.***

***(A)*** Hepatic $H_2S$, ***(B)*** Hepatic CSE gene expression, (***C***) Hepatic CSE protein expression, (***D***) Hepatic CSE enzyme activity, ***(E)*** Hepatic SQR gene expression, (***F***) Hepatic SQR protein expression. ***(G)*** Renal $H_2S$, ***(H)*** Renal CSE gene expression, (***I***) Renal CSE protein expression, (***J***)

Renal CSE enzyme activity, ***(K)*** Renal SQR gene expression, (***L***) Renal SQR protein expression. Data were analyzed with Student unpaired 2-tail *t* test. Data are presented as box plots (median, minimum, and maximum). CSE, cystathionine γ-lyase; SQR, sulfide quinone oxidoreductase.

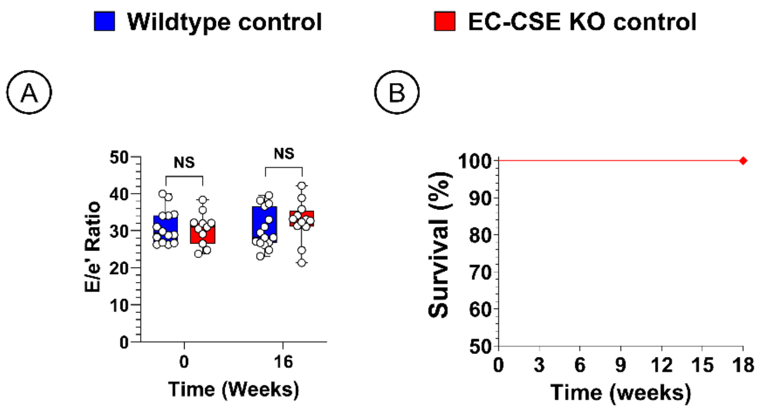


***Supplemental Figure 3. Cardiac Function and Survival Control Wildtype and EC-CSE KO Mice.***

(***A***) Ratio of early mitral diastolic inflow velocity (E) and mitral annular early diastolic velocity (e'), ***(B)*** Survival analysis., Data in panels ***A*** were analyzed with Repeated Measures Two-Way ANOVA. Survival data in panel ***B*** was analyzed with Kaplan-Meier survival analysis. Data in panels ***A and B*** are presented as box plots (median, minimum, and maximum).

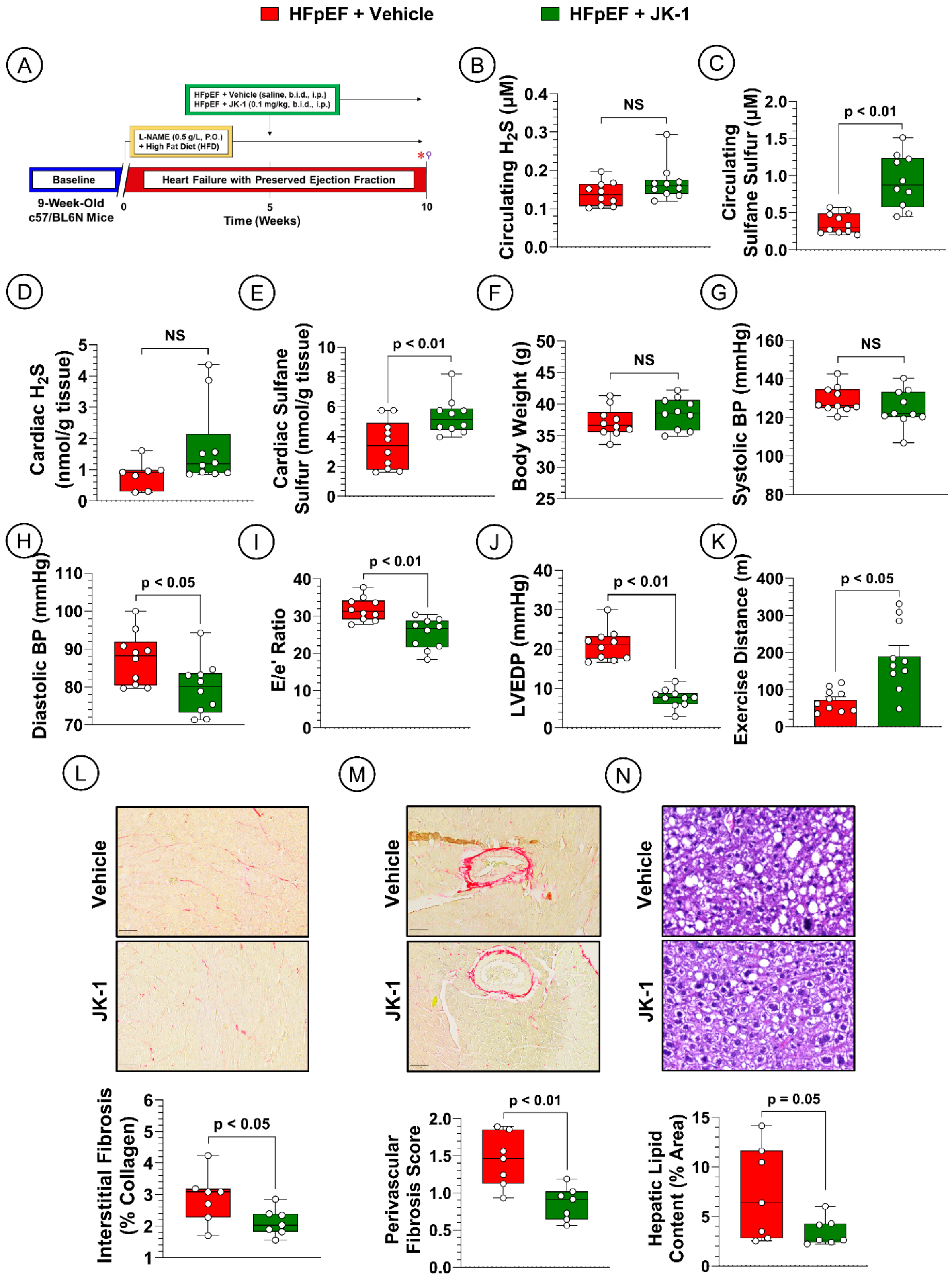
HFpEF + Vehicle
HFpEF + JK-1
A
HFpEF + Vehicle (saline, b.i.d., i.p.)
HFpEF + JK-1 (0.1 mg/kg, b.i.d., i.p.)
L-NAME (0.5 g/L, P.O.)
+ High Fat Diet (HFD)
Baseline
Heart Failure with Preserved Ejection Fraction
9-Week-Old
c57/BL6N Mice
Time (Weeks)
B
Circulating H2S (μM)
NS
C
Circulating Sulfane Sulfur (μM)
p < 0.01
D
Cardiac H2S (nmol/g tissue)
NS
E
Cardiac Sulfane Sulfur (nmol/g tissue)
p < 0.01
F
Body Weight (g)
NS
G
Systolic BP (mmHg)
NS
H
Diastolic BP (mmHg)
p < 0.05
I
E/e' Ratio
p < 0.01
J
LVEDP (mmHg)
p < 0.01
K
Exercise Distance (m)
p < 0.05
L
M
N
Vehicle
JK-1
Interstitial Fibrosis (% Collagen)
p < 0.05
Perivascular Fibrosis Score
p < 0.01
Hepatic Lipid Content (% Area)
p = 0.05

***Supplemental Figure 4. Hydrogen Sulfide Therapy Attenuates HFpEF Severity in Murine "Two-Hit" Model.***

(***A***) Study timeline, (***B***) Circulating $H_2S$, ***(C)*** Circulating sulfane sulfur, ***(D)*** Myocardial $H_2S$, ***(E)*** Myocardial sulfane sulfur, (***F***) Body weight, (***G***) Systolic blood pressure, (***H***) Diastolic blood pressure, ***(I)*** Ratio of early mitral diastolic inflow velocity (E) and mitral annular early diastolic velocity (e'), (***J***) Left ventricular end diastolic pressure (LVEDP), ***(K)*** Treadmill exercise distance, ***(L)*** Representative images of cardiac interstitial fibrosis in HFpEF (upper) and HFpEF + JK-1 (lower) animals stained with Masson's Trichrome and respective quantification, (***M***) Representative images of cardiac perivascular fibrosis stained with Masson's Trichrome and respective quantification, (***N***) Representative images of hepatic lipid accumulation stained with hematoxylin and eosin and respective quantification. Data were analyzed with Student unpaired 2-tail *t* test. Data in all panels are presented as box plots (median, minimum, and maximum) except data in panel ***K*** are presented as mean ± SEM. * Echocardiography and Treadmill exercise, ♀ Invasive hemodynamics, $H_2S$ and sulfane sulfur measurements, Cardiac fibrosis and hepatic lipid content.

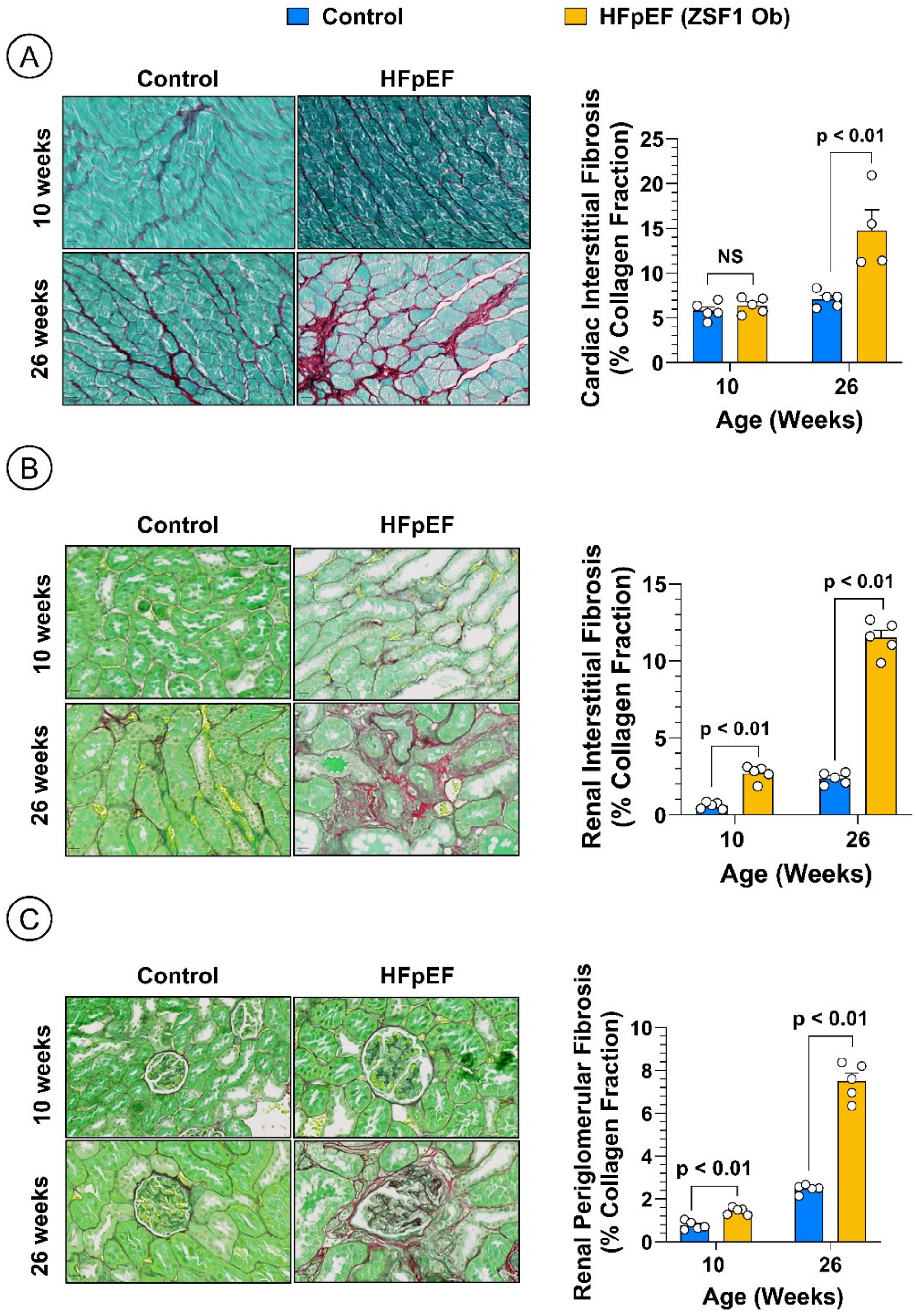


***Supplemental Figure 5. Fibrosis in the ZSF1 Obese Rat Tissues.***

(***A***) Representative images of cardiac interstitial fibrosis stained with Masson's Trichrome and fast green counterstaining and respective quantification, (***B***) Representative images of renal interstitial fibrosis stained with Masson's Trichrome and fast green counterstaining and respective quantification, (***C***) Representative images of renal periglomerular fibrosis stained with Masson's Trichrome and fast green counterstaining and respective quantification. Data were analyzed with multiple Student unpaired 2-tail *t* tests. Data are presented as mean ± SEM.

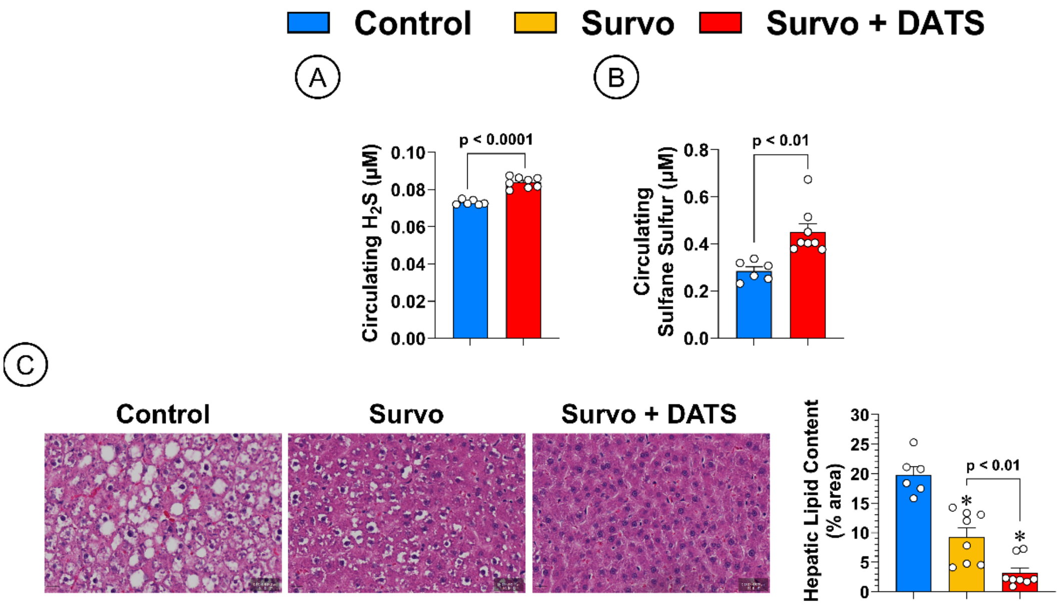


***Supplemental Figure 6. Circulating H2S and Hepatic Lipid Content in ZSF1 Obese Rats treated with Survodutide and DATS.***

(***A***) Circulating $H_2S$, (B) Circulating sulfane sulfur, (C) Representative images of hepatic lipid accumulation stained with hematoxylin and eosin and respective quantification. Data in panels ***A and B*** were analyzed with Student unpaired 2-tail *t* test. Data in panel C were analyzed with one-way ANOVA followed by Tukey test. Data are presented as mean ± SEM. * $p < 0.05$ vs. Control.
